# Fine-Mapping and Credible Set Construction using a Multi-population Joint Analysis of Marginal Summary Statistics from Genome-wide Association Studies

**DOI:** 10.1101/2022.12.22.521659

**Authors:** Jiayi Shen, Lai Jiang, Kan Wang, Anqi Wang, Fei Chen, Paul J. Newcombe, Christopher A. Haiman, David V. Conti

## Abstract

Recent advancement in Genome-wide Association Studies (GWAS) comes from not only increasingly larger sample sizes but also the shifted focus towards underrepresented populations. Multi-population GWAS may increase power to detect novel risk variants and improve fine-mapping resolution by leveraging evidence from diverse populations and accounting for the difference in linkage disequilibrium (LD) across ethnic groups. Here, we expand upon our previous approach for single-population fine-mapping through Joint Analysis of Marginal SNP Effects (JAM) to a multi-population analysis (mJAM). Under the assumption that true causal variants are common across studies, we implement a novel version of JAM that conditions on multiple SNPs while explicitly incorporating the different LD structures across populations. The mJAM framework can be used to first select index variants using the mJAM likelihood with any feature selection approach. In addition, we present a novel approach leveraging the ideas of mediation to construct credible sets for these index variants. Construction of such credible sets can be performed given any existing index variants. We illustrate the implementation of the mJAM likelihood through two implementations: mJAM-SuSiE (a Bayesian approach) and mJAM-Forward selection. Through simulation studies based on realistic effect sizes and levels of LD, we demonstrated that mJAM performs better than other existing multi-ethnic methods for constructing concise credible sets that include the underlying causal variants. In real data examples taken from the most recent multi-population prostate cancer GWAS, we showed several practical advantages of mJAM over other existing methods.

## Introduction

The development of high-throughput genotyping and genotype imputation has boosted the application of genome-wide association studies (GWAS) which is now a standard approach to identify susceptibility loci or genomic regions for many complex diseases and traits^1,2^. However, the linkage disequilibrium (LD) of single-nucleotide polymorphisms (SNPs) makes it challenging to determine the true causal variant(s) within a region or to further prioritize genetic variants for functional studies^2,3^.

Fine-mapping is a post-GWAS approach which seeks to specify the underlying causal variant and quantify the strength of effect given existing evidence that a certain region is likely to contain at least one causal signal. Many methods for fine-mapping often start with a lead SNP – the SNP with the smallest *p*-value within one region – and then they examine additional highly correlated neighboring SNPs in the region using different strategies such as setting a threshold on pairwise correlation (r^2^) with the lead SNP^2^. These approaches are intuitive but do not jointly analyze all the SNPs within a region. In addition, they often do not generalize easily to the investigation in multiple populations.

More recent and advanced fine-mapping approaches attempt to jointly or conditionally analyze all SNPs within a region, and include stepwise regression^2^, penalized regression^4–7^, and Bayesian methods^8–11^. Conditional step-wise selection has been used to discover multiple signals at a locus with individual level data^12,13^. However, stepwise selection can be very unstable with a large amount of highly correlated SNPs and the *P*-values of the signals in the final selected model tend to be conservative^2^.

Alternative selection approaches with individual level data are penalized regression models, such as lasso^4^ and elastic net^5^, and Bayesian methods, such as CAVIAR^10^ and Sum of Single Effect models (SuSiE)^11^. In contrast to step-wise selection, penalized regression techniques are potentially more stable because the penalty term encourages shrinkage of effect estimates towards zero resulting in sparsity and robust estimation. However, penalized often do not perform well with highly correlated SNPs and they do not represent the uncertainty in effect estimation and model selection^2^. In contrast, fully Bayesian methods compute posterior probabilities for models within the model space to infer the probability of causality for each SNP and often result in credible sets to measure the fine-mapping resolution using these probabilities. Ideally, exact inference is possible by enumerating all possible models or combinations of SNPs but the model space increases so rapidly that exhaustive searches become impractical as the number of SNPs increases. Stochastic search algorithms are often used to perform inference on posterior distributions. For example, piMASS^8^ and BVS^14,15^ both use Markov chain Monte Carlo (MCMC) algorithm to search through the model space, while the later can also incorporate external annotations as prior information in the Bayesian model selection to further prioritize causal SNPs.

In addition to analyses performed on individual-level data, methods for fine-mapping using only summary statistics from GWAS are becoming more widely applied ^16–19^. In general, these methods use reference samples to estimate the correlations between SNPs and then integrate the correlation structure with modified marginal SNP summary statistics in a multivariable regression framework to approximate the corresponding individual-level analysis. Differences between methods are due to variations in the assumptions for residual error and algorithms for model selection^16–19^. For example, FINEMAP^16^ places a Gaussian prior for causal effect estimates and adopts a shotgun stochastic search algorithm to prioritize the search to a set of most likely important causal configurations. The original implementation of the Joint Analysis of Marginal SNP Effects (JAM)^19^ invokes a Cholesky transformation on the linear regression likelihood, adopts a *g*-prior for effect estimates, and then implements a computationally efficient reversible jump MCMC stochastic search algorithm.

Leveraging the information across multiple ethnic groups or ancestry populations can enhance the power of fine-mapping^20–22^. Different ancestry groups may have distinct LD structures due to different evolutionary and migration histories^23,24^. For example, compared to non-African Americans, African Americans have smaller LD blocks with weaker correlations as the number of recombination events for each region is expected to be higher^25^. If a true causal variant exists across populations, its corresponding estimated association across populations should be more consistent than the estimated association for proxy SNPs with different LD across populations^26–28^. Therefore, integrating the difference in the LD structures across populations can potentially narrow the credible set that a causal variant resides in and improve the resolution of the fine-mapping^29,30^.

Here, we present an extension of the single-population fine-mapping through JAM to a multi-population setting by fitting a multi-SNP joint model, “mJAM”. mJAM assumes that the true causal variant(s) share the same effect across ancestry groups and it explicitly accounts for different LD structures across ancestry groups in the joint model. The mJAM likelihood allows for different feature selection procedures to be performed on summary statistics obtained from multiple populations. This includes Bayesian variable selection approaches that also yield credible sets or more conventional approaches for only selecting certain SNPs. When combined with approaches that only select specific SNPs, mJAM conditional models can further be used in a mediation type framework to construct credible sets for these index variants in a multi-population analysis. We illustrate this flexibility with two computationally efficient implementations of mJAM: “mJAM-SuSiE” for Bayesian variable selection with native SuSiE credible sets, and “mJAM-Forward” for frequentist forward selection of index SNPs and subsequent credible set construction. Through simulation studies with realistic effect size and various patterns of LD, we compare mJAM-SuSiE and mJAM-Forward with other multi-population approaches, including the most commonly used fixed-effect meta-analysis, COJO^17^ with pooled LD structure and meta-analyzed summary statistics, and MsCAVIAR^31^, a Bayesian fine-mapping approach that allows for an arbitrary number of causal variants in a region. We then applied these methods to three known regions for prostate cancer to demonstrate the practical advantages of mJAM.

## Material and methods

### Multi-population JAM

To simplify notation and without loss of generality, we consider the scenario with three populations. Within each population and for a given set of *p* SNPs within each region, a linear phenotypic model is used.

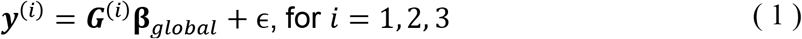

where ***y***^(*i*)^ is a *N*^(*i*)^ × 1 vector of mean-centered phenotypic trait values for the *i^th^* population, with *N*^(*i*)^ being the sample size of the *i^th^* population; ***G***^(*i*)^ is a *N*^(*i*)^-× *p* matrix of individual-level genotype data for the *i^th^* group, where each SNP has been centered to its mean; **β**_*global*_ ∈ ℝ^*P*^ denotes the joint effect of the given set of *p* SNPs. *ϵ* ~*N*(*O, σ*^2^) where *σ*^2^ is the residual variance. It is assumed that all three populations share the same joint effect size, i.e., **β**_*global*_, and the same residual variance.

Akin to a meta-regression, a second-stage model describes the relationship between the population joint effect estimates and the underlying true effect:

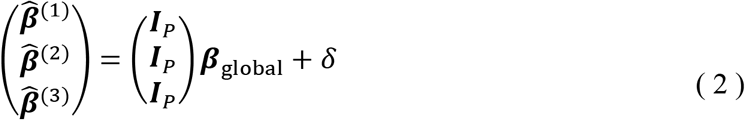

where *δ*~*N*(*0,τ*^2^) and 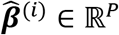 is the vector of estimated joint SNP effects for the *i^th^* population.

Equation (1) and (2) together form a two-stage model when individual-level data are available. The first stage is three separate linear phenotypic models whereas the second stage fits a fixed-effect meta-analysis model that combines all populations together. By replacing the ***β***_*global*_’s in (1) with (2), we have the following linear fixed-effect model that incorporates the individual-level data of all populations:

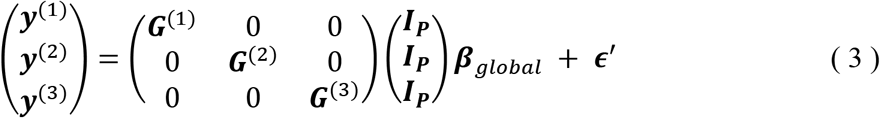

With summary data in which only the marginal effect sizes and their standard errors are available, it is also possible to estimate the joint effect size, ***β**_global_*, with an additional reference sample that estimates the LD between the SNPs^2,32^. Thus, Equation (3) can be used with only GWAS summary statistics with a modified mJAM likelihood after linear transformation:

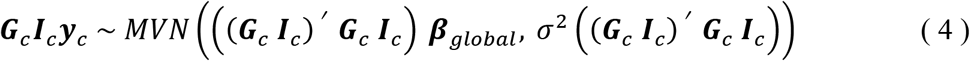

respectively. By expanding each matrix, we have 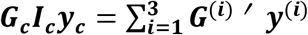 and 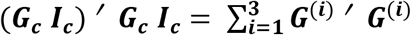 where **G**^*i*^’ ‘ ***G***^(*i*)^ and ***G***^(*i*)^’ ‘ ***y***^(*i*)^ are population-specific statistics and can be estimated by population-specific GWAS summary statistics and a reference genotype matrix or LD matrix. Detailed derivation can be found in Supplemental Methods.

### Index SNP Selection and Credible Set Construction for Fine Mapping

mJAM establishes a multi-SNP model within each population with corresponding population-specific LD, while jointly estimating a fixed-effects summary estimate of effect. The mJAM likelihood presented in Equation (4) can be used in a wide variety of existing feature selection approaches which are applicable to the mJAM statistics shown in Equation (4). Possible approaches for index SNP selection in mJAM includes stepwise selection^2^, Ridge regression^7^, and Bayesian approaches such as SuSiE^11^.

We adopt a forward selection approach based on conditional *P*-value for index SNP selection because of its computational efficiency and straightforward interpretation. We define our implementation of “mJAM-Forward” as a two-step approach in which a first step relies on a conventional stepwise forward selection to select an additional index SNP based on its corresponding *P*-value from a mJAM model conditional on any previous index SNP(s). We incorporate a *g*-prior to stabilize effect estimates^33^. To avoid fitting models with highly correlated SNPs we include a pruning process within Algorithm 1.

The second step for mJAM-Froward is to define a multi-population credible set for each index SNP. Here, we fit two mJAM models for each candidate credible set SNP, *W*, located within a region of an index SNP, *X*. These models demonstrate that the candidate credible set SNP is: 1) associated with disease marginally, and 2) that the index SNP mediates the effect of the candidate SNP on the disease. The first model includes *W* by itself to yield a probability that *W* is associated with the trait. This model also provides a posterior distribution for the marginal effect for the candidate credible set SNP.

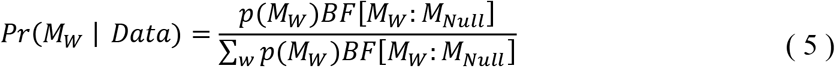

where *p*(*M_w_*) is the prior density of one-SNP model that includes *W* and *BF*[*M_W_: M_Null_*] is the Bayes factor of one-SNP model with *W* to the null model. See Supplemental Methods for detailed expression of *BF*[*M_W_: M_Null_*] with the incorporation of a g-prior of the effect estimates. The second model conditions on the index SNP, *X*, to obtain a posterior estimate for an adjusted effect estimate for the credible set SNP. Borrowing from a mediation framework^34^, we then calculate the probability that the index SNP mediates the candidate credible set SNP effect using the two models (Figure 1).

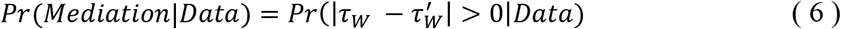

where *τ_w_* is the total effect of the candidate credible set SNP on the outcome and 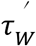 is the direct effect. A strong mediation effect indicates that the observed marginal effect of the candidate credible set SNP on the outcome is mainly due to its indirect effect through its strong correlation with the index SNP, and not due to a direct effect on the outcome. These two model probabilities are then combined to calculate the probability that a candidate SNP is a credible set SNP, Posterior Credible Set Probability (PCSP).

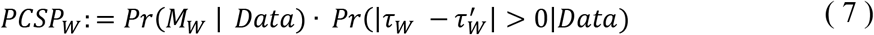

**Figure 1.**
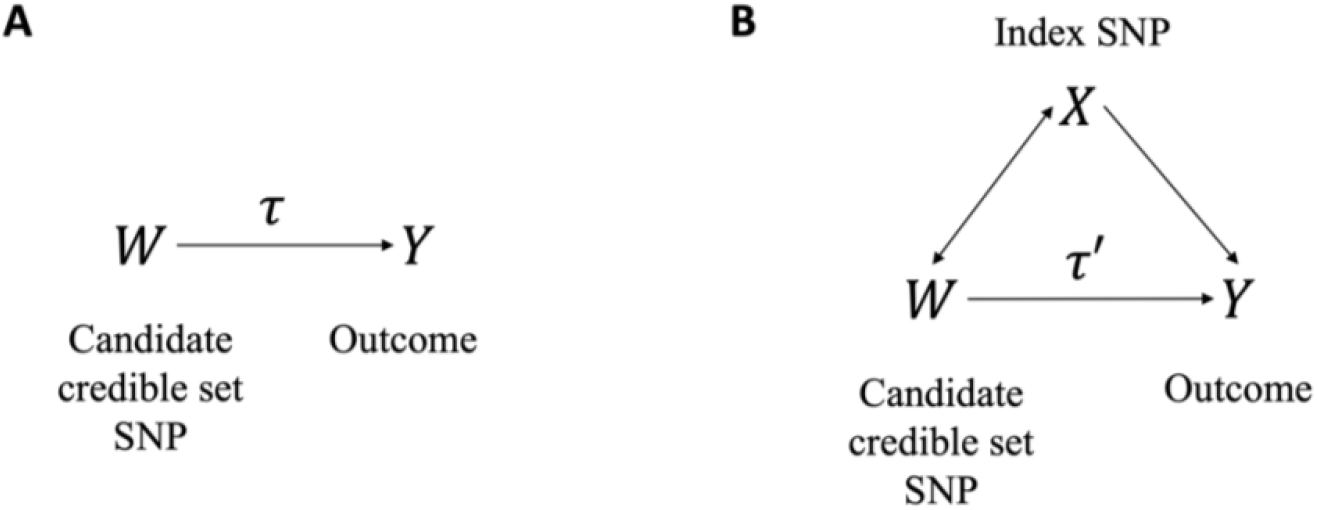
The direct acyclic graphs (DAG) for the probability that the index SNP mediates the candidate credible set SNP effect. (A) Model with the candidate credible set SNP, W, by itself. τ is the total effect of W on Y. (B) Model with W and X, the index SNP. τ’ is the direct effect of W on Y.

PCSP are then scaled over all SNPs in the region and used to define a 95% credible set of cross-population SNPs.

#### Algorithm 1 Pseudo algorithm for fitting mJAM-Forward and credible set construction in a region

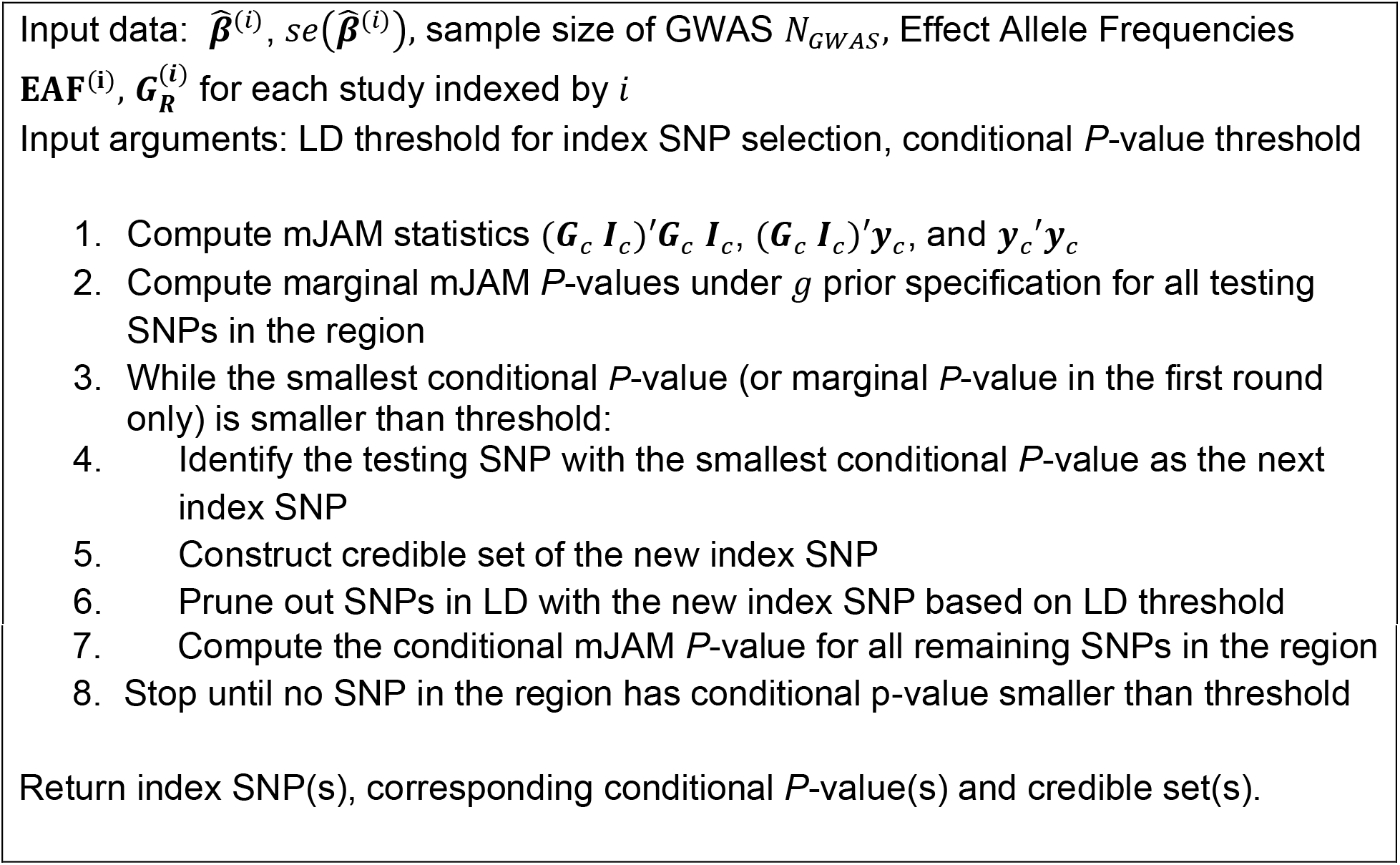

We also integrate the mJAM likelihood and summary statistics into a Bayesian selection method that indicates index SNPs and simultaneously estimates credible set SNPs, “mJAM-SuSiE” (See Supplemental Methods for the pseudo-algorithm of fitting mJAM-SuSiE)^35^.

### Incorporating missing variants in mJAM

In genetic association studies with more than one cohort or study, it is common that a particular SNP might be available in some studies but missing in the others^36^. A notable practical feature of the mJAM framework is that it allows for these SNPs with missing information to be analyzed without being filtered or removed. This is accomplished with a simple modification by substituting a value of zero in the identity matrix in Equation (3) and (4). Such modification then allows for observed statistics from other populations to be used but removes the contribution from the population in which it is missing but does not alter the algorithm nor the fitting process. This modification is applicable either when the SNP is missing in the reference panel or when the population-specific GWAS summary statistics are not available for the SNP (See Supplemental Methods for more details).

### Simulation Study on Structured LD

We conducted a simulation study to compare the performance of the two mJAM implementations (mJAM-SuSiE and mJAM-Forward) with three commonly used alternative approaches: fixed-effect meta-analysis, COJO stepwise selection and MsCAVIAR. Fixed-effect meta-analysis takes an inverse-variance weighted average of the marginal estimates from individual studies or populations. COJO approximates the conditional and joint effect from summary statistics and single reference LD and then implements a stepwise selection based on conditional *P*-values. Additionally, for use of COJO on multiple populations, the summary-level statistics come from the fixed-effect meta-analysis across all populations and the reference LD can be obtained from either the pooled individual-level genotype data or a subset of meta-analysis sample. We used the former as the reference LD for COJO in our simulations. MsCAVIAR is built upon a Bayesian multivariate normal framework first described as CAVIAR^10^ to account for between-study or between-population heterogeneity using a random-effects model. To compare the performance of index SNP selection across these multi-population approaches, we use three metrics: number of selected index SNPs, sensitivity/power, and positive predictive value (PPV). In addition, since MsCAVIAR, mJAM-SuSiE, and mJAM-Forward output credible set(s) within each region, we compare credible set performance using the number of credible set(s), size of each credible set, sensitivity/power, PPV and empirical coverage. For the two non-Bayesian methods, FE and COJO, we consider the group of SNPs with meta-analyzed or conditional *P*-values less than a Bonferroni-corrected significance level as a single credible set for the purpose of performance comparison.

We performed two sets of scenarios: 1) simulated correlation structures with the same block LD structures across populations; and 2) simulated correlation based on real genetic correlation structures observed in the study cohort from Elucidating Loci Involved in Prostate Cancer Susceptibility (ELLIPSE) OncoArray Consortium^21^. For the first set of scenarios, we first simulated a baseline scenario where each population has 3 individual association studies with N = 5,000 each to closely represent the real-life situation where there are multiple association studies performed for each ethnic group (total sample size = 5,000 × 3 studies/population × 3 population = 45,000). A total of 50 SNPs are simulated in 5 blocks of 10 SNPs. Within each block of 10 SNPs, the pairwise correlations are uniformly set to a constant value r^2^ across ancestries for simplicity. r^2^ varies from 0, 0.6^2^ and 0.9^2^ to represent independent, moderate LD and high LD scenarios. Corresponding LD heatmaps are shown in Figure S1. We then selected a single causal SNP with an effect size of 0.03 for a standard normal outcome. The baseline scenario was extended by varying parameters, including the ratio of sample sizes between each population, levels of LD, the total number of causal SNPs and corresponding effect sizes.

### Simulation Study with Real Data

To better capture realistic LD patterns, we performed simulations based on real correlation within three ancestry groups (Europeans, African Americans, and East Asians) from the ELLIPSE OncoArray Consortium^21^. The available sample sizes for these three populations are 93,749 Europeans, 9,531 African Americans, and 2,075 Asians. We simulated 120 SNPs within a 1334 kb region from chromosome 2 using a multivariate normal model with an estimated correlation structure from individual-level genotypes. The heatmap of this region for each ethnicity is shown in Figure S2. In each simulation, we randomly chose one SNP out of a selected LD block to be the causal SNP with effect size being 0.04, resulting in an empirical average -log10(*P*-value) of the most significance variant of 7.75 (*P*-value ≈ 1.8 × 10 ^−8^) averaged across 500 simulations.

### Applied examples

To illustrate mJAM on real data, we applied the methods on three regions using summary statistics from the latest cross-ancestry prostate cancer association study^37^ across four ancestry groups, including 122,188 prostate cancer cases and 604,640 controls of European ancestry, 19,391 cases and 61,608 controls of African ancestry, 10,809 cases and 95,790 controls of East Asian ancestry, and 3,931 cases and 26,405 controls from Hispanic populations. Within each region, we applied mJAM-Forward to select index SNP(s) using population-specific summary statistics and reference dosage for each population. Then we constructed mJAM credible set(s) by including top SNPs ranked by their mJAM posterior probabilities until those SNPs included in the credible set reached a cumulative posterior probability of 95%. Reference dosage were obtained from the Prostate Cancer Association Group to Investigate Cancer-Associated Alterations in the Genome and Collaborative Oncological Gene-Environment Study Consortium [PRACTICAL iCOGS], the Elucidating Loci Involved in Prostate Cancer Susceptibility OncoArray Consortium [ELLIPSE OncoArray], the African Ancestry Prostate Cancer Consortium [AAPC GWAS], GWAS of prostate cancer in Latinos [LAPC GWAS] and Japanese [JAPC GWAS]^21^. Results from mJAM-Forward are compared with those from mJAM-SuSiE, COJO and MsCAVIAR.

## Results

### Simulation Study on Artificial LD

Under the baseline scenario (50 SNPs in total, 1 causal SNP with an effect size of 0.03, 3 studies per population, and balanced sample size across populations), the 95% credible sets from mJAM-Forward, mJAM-SuSiE, and MsCAVIAR were well calibrated to the specified coverage level (Figure S3). Both mJAM-Forward and mJAM-SuSiE preserved relatively high sensitivity in terms of including the true causal SNP in its credible set (sensitivity = 0.86 and 0.64 respectively, Figure 2A). Although MsCAVIAR had the highest sensitivity (0.99) under the baseline scenario, its average credible set size was much larger (9.47 for MsCAVIAR; 2.12 for mJAM-Forward and 0.78 for mJAM-SuSiE, Figure 2C), thus leading to a much lower PPV (0.39, Figure 2B). mJAM-SuSiE had the highest PPV (0.89) among the methods we compared, meaning that it had the highest proportion of true causal over the total number of credible set SNPs on average, followed by mJAM-Forward (0.58) (Figure 2B). In terms of credible set sensitivity, PPV and average CS size the methods had similar patterns of performance for scenarios expanded beyond the baseline to various LD structures, imbalanced sample size across populations, and 3 causal SNPs (Figure S4).

**Figure 2.**
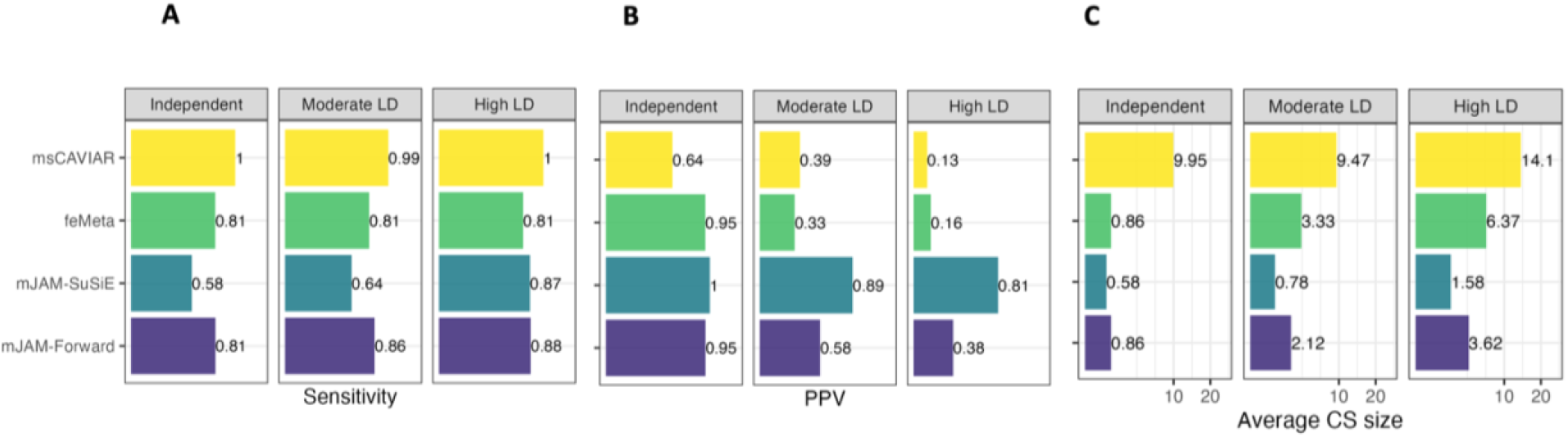
Credible set performance in simulation studies with artificial LD structure. (A) Sensitivity, i.e. the proportion of 500 simulations where the true causal SNP was selected in a credible set. (B) Positive Predictive Value (PPV), i.e., the proportion of true causal SNP over the credible set size, averaged over 500 iterations. (C) Average CS size.

In terms of identifying the true causal variant as an index SNP (i.e. sensitivity), mJAM-Forward and MsCAVIAR had the best performance under moderate LD scenarios (Figure 3A) with a sensitivity was 0.73, and 0.72 respectively. However, for these two methods, mJAM-Forward had a better PPV was 0.81, compared to MsCAVIAR (0.72). In comparison, mJAM-SuSiE had poor sensitivity (0.62) but a higher PPV (0.88). COJO had a similar performance with mJAM-Forward under independent LD scenarios but its sensitivity and PPV worsened compared to mJAM-Forward as the level of LD increases. Though COJO performs a similar stepwise selection as mJAM-Forward, unlike mJAM-Forward that specifically accounted for population-specific LD, COJO uses meta-analyzed marginal summary statistics and pooled LD panel which makes it difficult to identify the true common variants through disentangling the population-specific LD structure. All methods selected on average 1 index SNP among 500 simulations, close to the true number of causals (Figure 3C). For MsCAVIAR pre-specification is required so we set the value to 1 for all scenarios. Notably for practical implementation, for a small number of scenarios (40%), mJAM-SuSiE did not select any index SNP under independent or moderate LD scenarios, leading to relatively low sensitivity compared other methods when averaged over replicates (Figure 3A).

**Figure 3.**
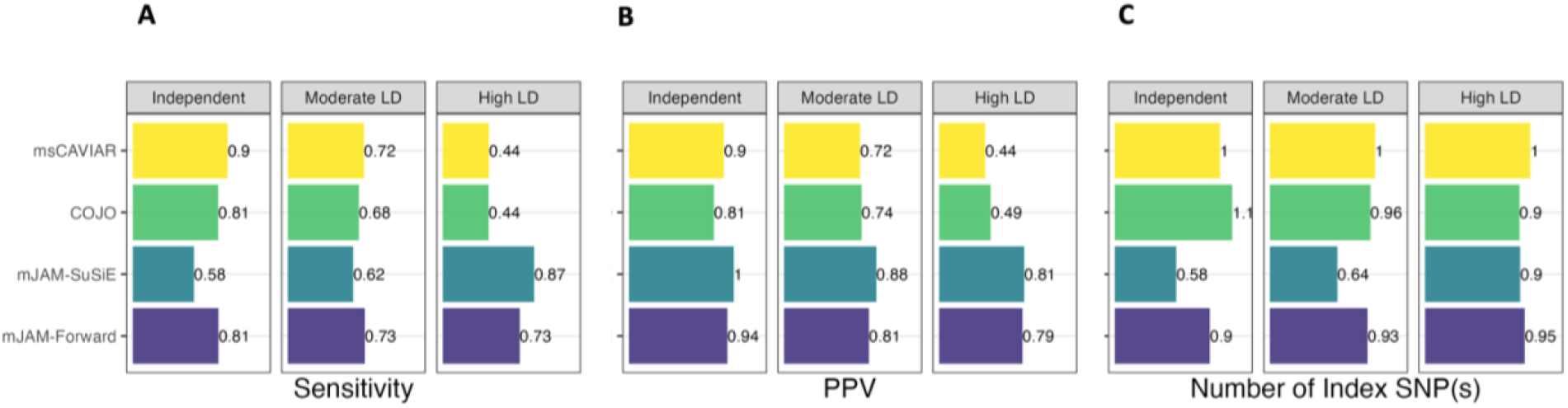
Performance of index SNP(s) selection in simulation studies with artificial LD structure. (A) Sensitivity, i.e. the proportion of 500 simulations where the true causal SNP was selected in an index SNP. (B) Positive Predictive Value (PPV), i.e., the proportion of causal SNP selected as an index over all selected indices, averaged over 500 iterations. (C) Number of index SNP(s) selected, averaged over 500 iterations.

When the pairwise correlation within each LD block increased, the average credible set sizes for all methods increased correspondingly (Figure 2C). As a result, under high LD scenarios, the PPV of identifying the true causal(s) in a credible set decreased to a noticeable extent for MsCAVIAR, mJAM-Forward, and FE (Figure 2B). Though mJAM-Forward’s PPV dropped due to the increase in credible set sizes on average, mJAM-Forward was still able to retain a sensitivity of 0.88 under the high LD scenario. mJAM-SuSiE achieved the highest PPV (0.81, Figure 2B) among all methods under high LD scenarios while retaining relatively high sensitivity and small credible set size. However, mJAM-SuSiE’s sensitivity was relatively low compared to mJAM-Forward and MsCAVIAR under independent or moderate LD scenarios (Figure 2A).

Despite of mJAM-SuSiE’s outstanding performance under high LD scenarios with moderate causal effect size, we noticed that its results were very sensitive to the marginal significance of the true causal SNPs. To represent a real-life situation where a lead variant within a region has an extremely significant marginal *P*-value, we expanded the baseline scenario with 1 true causal SNP to additional scenarios with increasing significance of the true causal SNP, where the average -log10(*P*-value) of the true causal ranges from 5 to 263 (mimicking significance often found in applied GWAS). Under increasingly high-power scenarios, mJAM-Forward consistently selected 1 credible set regardless of the significance of the true causal whereas the average number of credible sets by mJAM-SuSiE increased as the statistical significance (i.e. effective power) increased (Figure 4B). As a result, mJAM-SuSiE selected more false positive SNPs within the credible sets when the true causal SNP has high observed marginal significance. In addition, the empirical coverage of mJAM-SuSiE’s credible sets dropped below the expected level quickly after the true causal SNP became more significant (Figure 4A). In contrast, mJAM-Forward’s credible sets remained well-calibrated.

**Figure 4.**
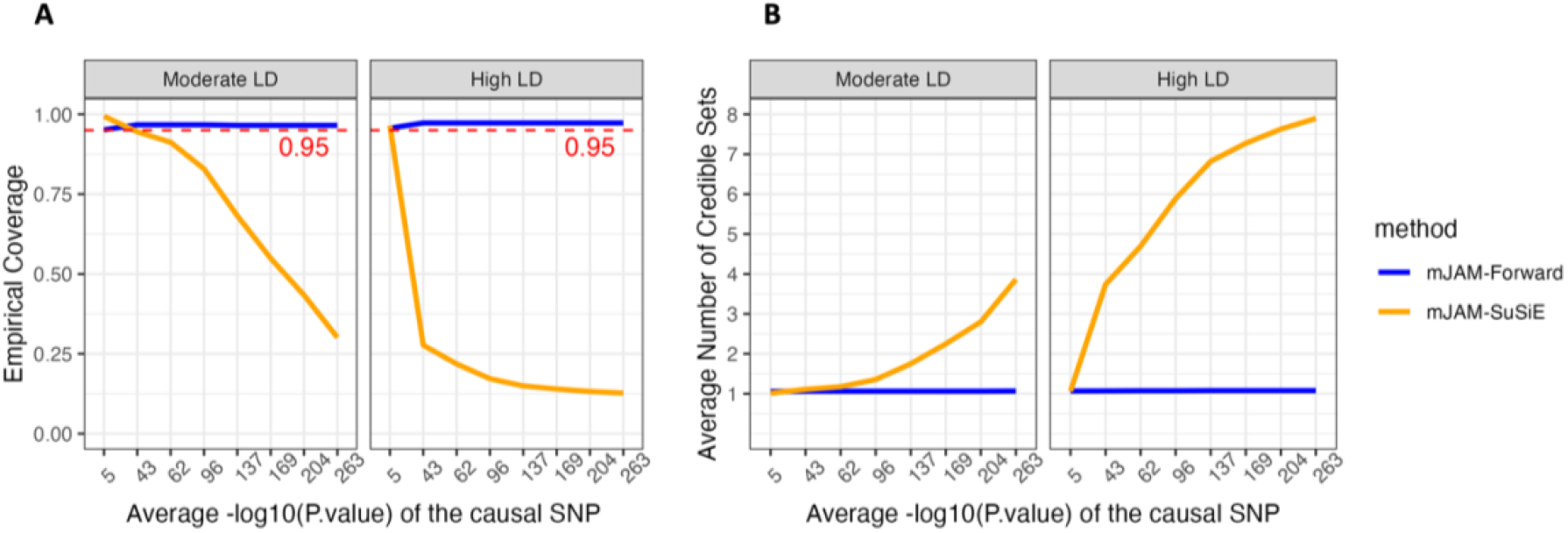
Credible set behaviour of mJAM-SuSiE and mJAM-Forward as causal SNP significance increases. Simulations were conducted under baseline scenario setting (1 causal SNP out of 50 SNPs in total which are divided into 5 LD blocks) with varying effect sizes. The average empirical -log10(P-value) of the causal SNP ranged from 5 to 263, covering most situations seen in practice. Red dashed line indicates requested coverage which is set to be 0.95 for both methods. (A) Empirical credible set coverage; (B) Average number of credible sets selected among 500 simulations.

To explore the impact of two types of missingness on the performance of mJAM-Forward, we modified our simulation studies with artificial LD structure to include a missing SNP in LD with the causal SNP, or with the missing SNP as the causal SNP itself. The flexibility of mJAM likelihood (Equation 2) allows us to incorporate SNPs with missing information in some studies or populations in the analysis. We found that when the missing SNP is in LD with the causal SNP, mJAM-Forward has stable performance in comparison to when there is no missingness (Figure S6). When the causal SNP is missing, mJAM-Forward still preserves the power both to select the causal SNP as the index SNP and to include the causal SNP in its credible set.

### Simulation Study on Real LD

When applied to the simulated data on the 120-SNP region on chromosome 2, mJAM-Forward, mJAM-SuSiE and MsCAVIAR selected on average around 1 index SNP whereas COJO selected 1.5 index SNPs, indicating a slight increase in false positive signals. mJAM-Forward had highest sensitivity and PPV of identifying the true causal from a complicated LD structure as an index SNP (Table 1). In terms of credible set performance, MsCAVIAR demonstrated high empirical coverage of its credible set as well as high sensitivity compared to the other two mJAM methods. However, such high sensitivity and PPV was achieved at the cost of a much larger size for the credible sets. The average size of the 95% CS of MsCAVIAR is 56.52, even larger than the number of SNPs that reached marginal genome-wide significant (5 × 10^-8^) in a fixed-effect meta-analysis (48.88). On the other hand, the average credible set size for mJAM-Forward and mJAM-SuSiE was 19.70 and 18.37 respectively. Meanwhile, both approaches preserved reasonably high sensitivity and empirical coverage.

**Table 1.** Comparison of model performance on data simulated from real LD structure.

|  |  | Method |  |  |  |  |
| --- | --- | --- | --- | --- | --- | --- |
|  |  | mJAM-Forward | mJAM-SuSiE | FE | COJO | MsCAVIAR |
| Credible Set Performance | Sensitivity<sup>a</sup> | 0.930 | 0.910 | 0.972 | - | 0.994 |
|  | PPV<sup>b</sup> | 0.064 | 0.069 | 0.024 | - | 0.022 |
|  | CS size<sup>c</sup> | 19.70 | 18.37 | 48.88 | - | 56.62 |
|  | CS coverage<sup>d</sup> | 0.934 | 0.940 | - | - | 1.000 |
| Index SNP Performance | Sensitivity<sup>e</sup> | 0.218 | 0.174 | - | 0.186 | 0.134 |
|  | PPV<sup>f</sup> | 0.219 | 0.021 | - | 0.144 | 0.134 |
|  | Number of selected index | 1.00 | 0.97 | - | 1.51 | 1.00 |

Abbreviations: FE, fixed-effect meta-analysis; CS, credible set, PPV, positive predictive value.
<sup>a</sup> proportion of true causal SNPs being selected in a credible set, averaged over 500 simulations
<sup>b</sup> proportion of true causal SNPs over the total number of selected credible set SNPs, averaged over 500 simulations.
<sup>c</sup> total number of SNPs included in a credible set, averaged over all 95% credible sets in 500 simulations.
<sup>d</sup> proportion of 95% credible sets in 500 simulations that included at least one true causal SNP.
<sup>e</sup> proportion of true causal SNPs being selected as an index SNP, averaged over 500 simulations.
<sup>f</sup> proportion of true causal SNPs over the total number of selected index SNPs, averaged over 500 simulations.

### Applied example 1: a single-hit region on chromosome 12

The first applied example is a 1013 kb region on chromosome 12 which consists of 276 SNPs with a marginal meta-analyzed *P*-value < 10^-3^ and minor allele frequency (MAF) > 2%. Figure S7 shows the LD structure for the four ancestry groups in this analysis. None of the SNPs in this region reached genome-wide significance in any population-specific analyses (Figure 5B) but after multi-population meta-analysis 48 SNPs are genome-wide significant (Figure 5A). By setting a conditional *P*-value threshold at 5 × 10^-8^, mJAM-Forward identified one index SNP at 12:109994870:A:T (meta P-value = 3.5 × 10^-10^) with a corresponding 95% credible set of 41 SNPs. The median r^2^ between the credible set SNPs with the index SNP is 0.998 for European LD, 0.979 for African, 0.990 for Hispanic and 0.996 for East Asian. COJO identified the same index SNP, 12:109994870:A:T. MsCAVIAR reported a slightly larger 95% credible set than mJAM-Forward, consisting of 45 SNPs (Figure S8). The index SNP of MsCAVIAR’s credible set is 12:109998097:A:G (meta *P*-value = 3.7 × 10^-10^) whose r^2^ with 12:109994870:A:T is greater than 0.99 in all four ancestry groups. This index SNP, 12:109998097:A:G, is included in a mJAM-Forward credible set only when coverage is increased to 99%; whereas the index SNP for mJAM-Forward, 12:109994870:A:T, is included in the 95% MsCAVIAR credible set. mJAM-SuSiE estimates a single 95% credible set with 28 total SNPs and a unique single index SNP, 12:109996343:A:C (meta *P*-value = 2.2 × 10^-9^) which is also included in both credible sets of mJAM-Forward and MsCAVIAR. The median r^2^ within a credible set is also greater than 0.99 for all ancestry groups (Table S1). The index SNP from mJAM-Forward was also included in its credible set (Figure S8B).

**Figure 5.**
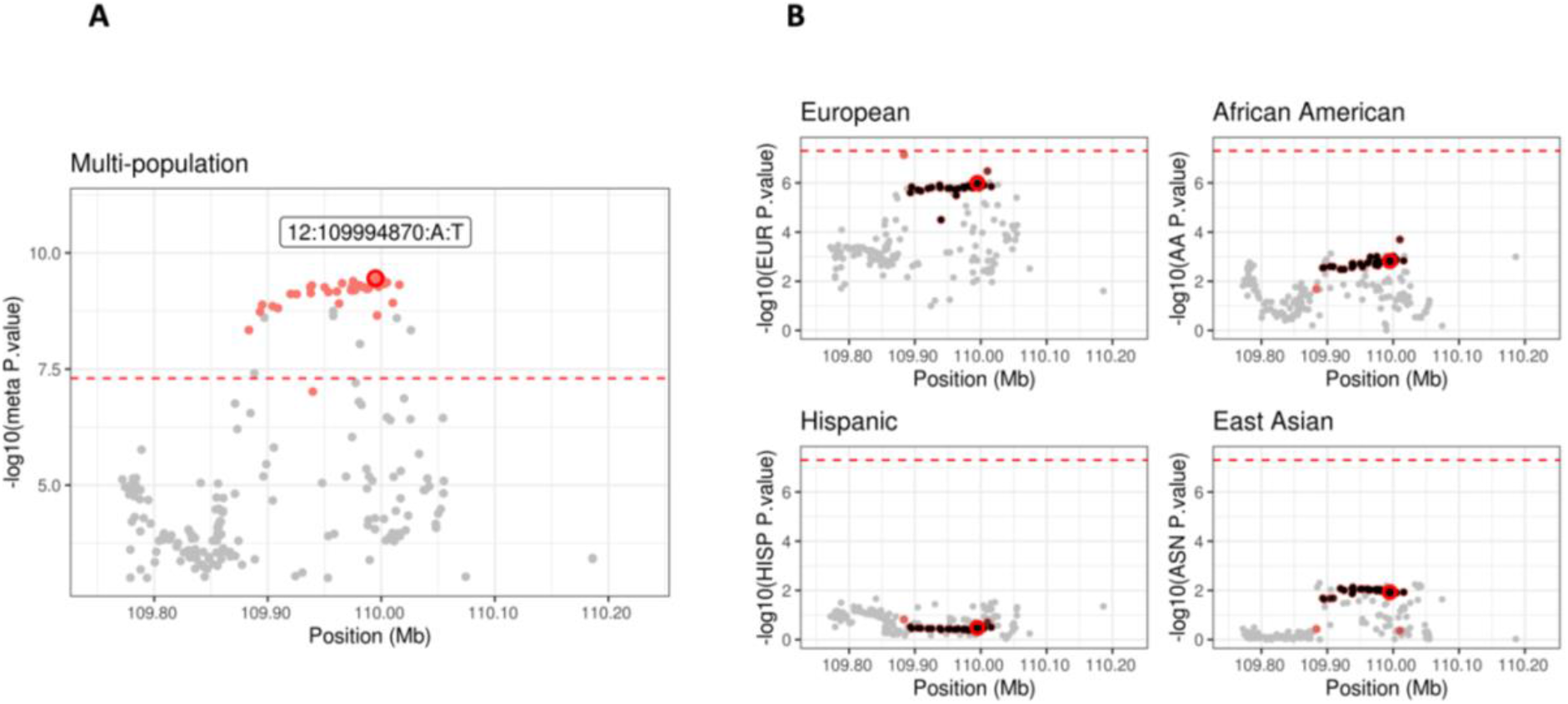
Manhattan plot for mJAM-Forward credible sets at chromosome 12 position 109194870 to 110794870. (A) y-axis is meta-analyzed -log10(P-value) from multi-ethnic analysis; (B) y-axis is -log10(P-value) from ethnic-specific analysis.

### Applied example 2: Asian-driven signals on chromosome 10

As a second example, we conducted an analysis on a chromosome 10 region which consists of 412 SNPs after QC and spans around 1571 kb. Figure S9 shows the LD structure in this region separately for European, African, East Asian, and Hispanic populations. This region contains two clear signals with meta-analyzed *P*-value < 10^-15^, which are mainly driven by the results from East Asian and African populations (Figure 6). In this example, mJAM-Forward identified two index SNPs, 10:80835998:C:T (meta *P*-value = 9 × 10^-21^ and 10:80238015:C:T (meta *P*-value = 1 × 10^-19^) (Figure 6A). The 95% mJAM-Forward credible set for the first index SNP, 10:80835998:C:T, contains 3 SNPs in total and there are 45 SNPs in the credible set for the second index SNP. The minimum r^2^ between the mJAM-Forward credible set SNPs with its own index SNP is no less than 0.95 in European, East Asian and Hispanic populations, and no less than 0.81 in African ancestry populations (Table S2). COJO identified two index SNPs, 10:80835998:C:T and 10:80240493:A:G. 10:80835998:C:T is the same as one of the index SNPs selected by mJAM-Forward and 10:80240493:A:G is included in the mJAM-Forward 95% credible set of 10:80238015:C:T. Since MsCAVIAR does not support reporting more than one distinctive credible set, we split this region into two adjacent regions and applied MaCAVIAR on these two subregions separately. MsCAVIAR selected the same 3-SNP 95% credible set (Figure S10) with index SNP being 10:80835998:C:T, and another 45-SNP credible set with index SNP being 10:80238015:C:T where 42 of them are replicated in the mJAM-Forward credible set. mJAM-SuSiE also identified the same 3-SNP credible set (95%) with the same index SNP 10:80835998:C:T but did not identify any credible set around 10:80238015:C:T. Instead, it reported two additional credible sets at 10:80260938:V1 (meta *P*-value = 2 × 10^-10^) and 10:80476778:V1 (meta *P*-value = 4 × 10^-4^) (Figure S10), and the credible set size is 2 and 5 respectively.

**Figure 6.**
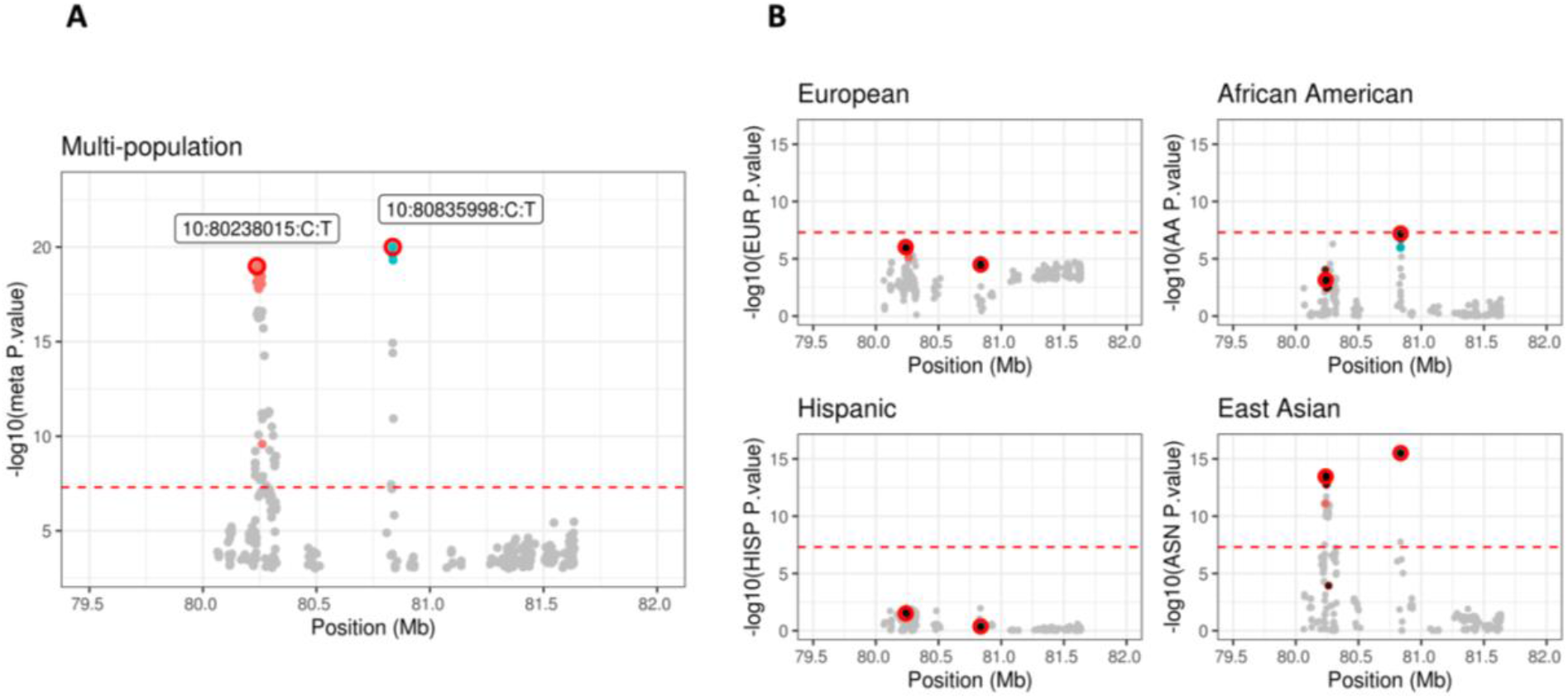
Manhattan plot for mJAM-Forward credible sets at chromosome 10 position 79436999 to 81635998. (A) y-axis is meta-analyzed -log10(P-value) from multi-ethnic analysis; (B) y-axis is -log10(P-value) from ethnic-specific analysis.

### Applied example 3: Secondary signal within 40kb region of a leading SNP

The third applied example illustrates a scenario where there is a secondary signal within close proximity of the leading SNP in a chromosome 11 region. This region spans 335.5 kb and consists of 191 SNPs that passed QC. The population-specific LD structure and Manhattan plot of multi-population meta-analysis results are shown in Figure S11 and Figure 7. The lead variant, 11:102401661:C:T, has a multi-population meta-analyzed *P*-value of **1.5** × **10^-38^** and mJAM-Forward identified a secondary index SNP, 11:102440927:A:G, only 39 kb away which has a meta *P*-value of **4.9** × **10^-11^**. The r^2^ between these two index SNPs is less than 0.01 in all four ancestry groups (Figure S11), suggesting statistical independence between these two SNPs. COJO selected the same primary index SNP, 11:102401661:C:T, and a different secondary index, 11:102433309:A:G, which has a meta *P*-value of 1.3 × 10^-7^ and is highly correlated with 11:102440927:A:G (r^2^ = 0.79 in EUR; 0.55 in AA; 0.87 in LA and 0.99 in ASN). mJAM-SuSiE also selected two credible sets in this region: the first set has 2 SNPs which are both replicated in mJAM-Forward’s first credible set; the second set has 26 SNPs where 24 of them are found in mJAM-Forward’s second set. However, the index SNP of the second set in mJAM-SuSiE is one with lower marginal significance (meta *P*-value = **6.3** × **10^-5^**) compared to mJAM-Forward.

**Figure 7.**
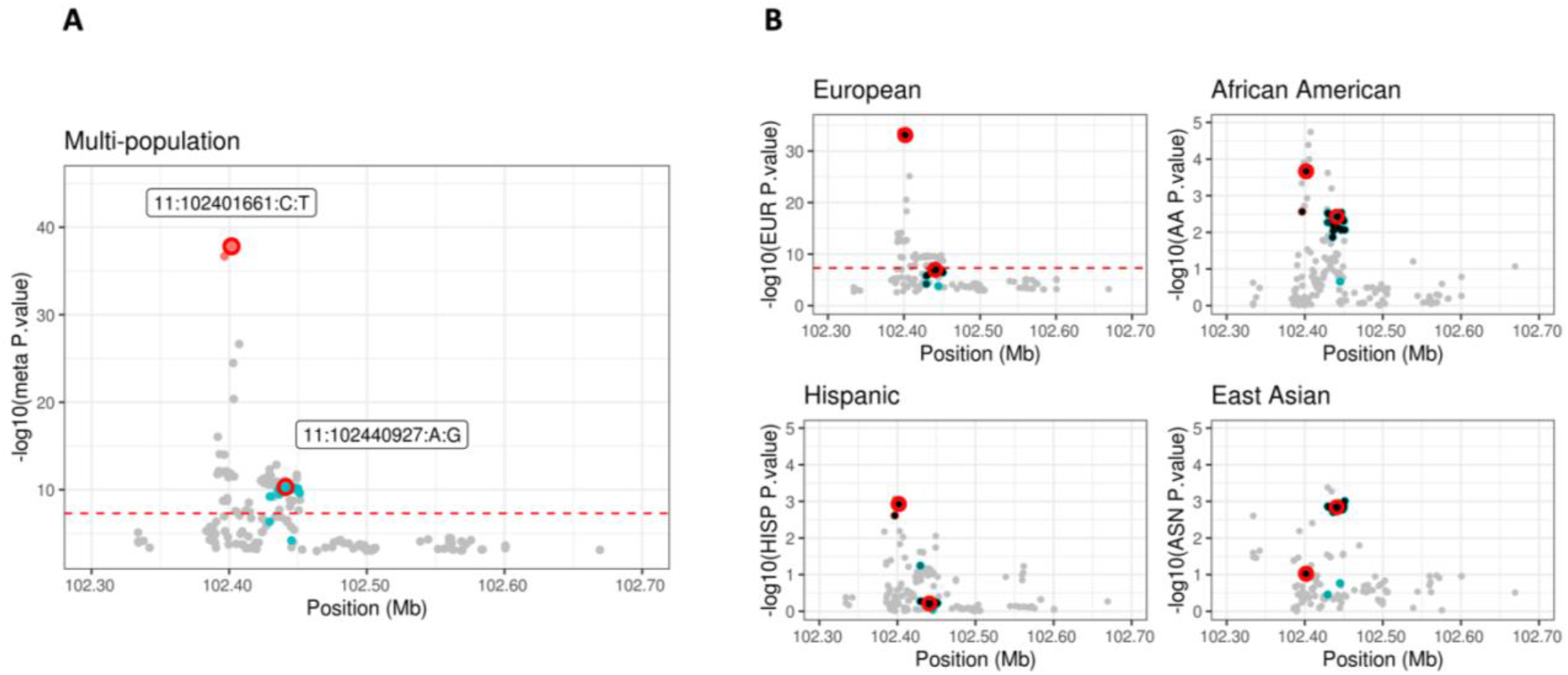
Manhattan plot for mJAM-Forward credible sets SNPs at chromosome 11 position 101601661 to 103201661. (A) y-axis is meta-analyzed -log10(P-value) from multi-ethnic analysis; (B) y-axis is -log10(P-value) from ethnic-specific analysis.

Both mJAM-SuSiE and mJAM-Forward are able to identify multiple sets within one region without any pre-defined number of causal variants. On the other hand, the implementation of MsCAVIAR requires users to specify the maximum number of causal variants in a region to enumerate all possible causal configurations. Gauging the possible number of causal variants can be difficult when secondary signals are located close to the lead variant. In this example, the secondary signal is located only 39 kb away from the leading variant, and visual inspection of the Manhattan plot (Figure 7) suggests only one peak. Even if we specify the number of causal variants to be two when applying MsCAVIAR to this region, MsCAVIAR reports only one credible set such that the posterior probability of this set containing 2 causal variants is at least 0.95. Thus, it becomes difficult to separate the selected credible set SNPs into two distinctive groups. When the number of causal variants is set to two, MsCAVIAR selected 24 SNPs among which the 2 SNPs with highest posterior probability are 11:102401661:C:T and 11:102396607:C:T (Figure S12). However, these two SNPs are in high LD and thus are likely linked to a single underlying causal signal and not indicative of multiple independent signals.

## Discussion

As integrating studies from ancestrally diverse populations may increase power to detect novel variant and improve fine-mapping resolution^22,38,39^, we extend our previous single-population fine-mapping through JAM to a multi-population approach, mJAM. mJAM requires only population-specific summary statistics and population-specific reference LD panels, which are more accessible than individual-level data to many researchers. mJAM explicitly incorporates the different LD structures across populations to yield conditional estimates of SNP effects from a single joint model. The mJAM framework can be used to first select index SNPs using existing feature selection approaches, such as forward stepwise selection^2^, Bayesian model selection^8,9,11^, or regularized regression^6,7^. To demonstrate this flexibility, we have implemented mJAM through two implementations of feature selection: mJAM-SuSiE (a Bayesian approach) and mJAM-forward selection. We also combine the forward selection implementation with a second step to identify credible set SNPs. This step works given any set of index SNPs within a region by estimating a posterior credible set probability (PCSP) for a SNP defined as a combination of two component probabilities: one models the marginal association between the candidate SNP and the outcome; the other models the mediation effect of the index SNP on the candidate SNP, borrowing from a mediation framework. These PCSPs are then used to construct credible sets. The closed-formed expression for PCSP allows computational efficient construction of credible sets, compared to other Bayesian approaches that often use computationally intensive algorithms to obtain posterior distributions. It also allows credible set construction from any index SNP list allowing researchers to apply other feature selection methods or use existing lists or knowledge to determine index SNP.

The two-stage model framework utilized in mJAM builds upon previous work highlighting the use of hierarchical JAM (hJAM)^40^, an approach for the joint analysis of marginal summary statistics that incorporates a prior information matrix. This matrix characterizes the SNPs and can include information such as SNP effects on gene expression analogous to TWAS or on intermediates biomarkers analogous to Mendelian randomization. mJAM is an extension to hJAM in that it replaces the prior information matrix in hJAM with a stacked identity matrix, 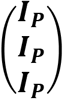, as described in Methods section. The stacked identify matrix can be interpreted as our prior believe on the joint SNP effect estimates that all populations share the same true effect sizes.

In a set of realistic simulation settings, both mJAM implementations demonstrated the ability to infer the number of independent signals within a region, to differentiate signals from noise, and to achieve a sufficient level of sensitivity while preserving high fine-mapping resolution through small-sized credible sets. We also investigated the impact of imbalanced sample size across populations on model performance and demonstrated that all methods showed a similar decrease in terms of sensitivity and PPV when the sample size is imbalanced but the total sample size remains constant (Figure S4). mJAM is described using three populations in simulation studies and we apply mJAM to real data with four distinct populations. In practice, mJAM can be used to analyze a large number of studies or population-specific summary statistics facilitating flexibility in application. Thus, analyses do not need to be limited to aggregating continental ancestry populations, but can include numerous, more specific ancestry appropriate reference panels to aggregate data across many studies (Figure S5). However, as with all summary statistic approaches that rely on reference panels, the ability to disentangle highly correlated SNPs will be driven by the sample sizes^41^ and LD within and between the reference panels used^42^. In addition, another practical limitation to many summary statistics-based approaches is the requirement for complete summary statistics and refence data for all SNPs across all studies and populations analyzed^36^. Missingness can be due to the difference in genotyping arrays used by different studies, or rare variants not being captured due to limited sample size in certain studies. Filtering too many variants might be dangerous because as less information is used to disentangle the LD structure within each region and potentially missing the causal variant. An important feature of mJAM is that it will work even in the presence of differential missingness across studies or populations utilizing all information that is available.

In the simulation study with artificial LD structures, mJAM-SuSiE resulted in outstanding performance under high LD scenarios, achieving both high sensitivity and high PPV. However, as the significance of the causal variant(s) within a region increases, mJAM-SuSiE tends to break down selecting more false positive signals with each in separate credible sets. This results in a substantial decrease in the empirical coverage of mJAM-SuSiE credible sets. In practice, we recommend limiting the application of mJAM-SuSiE to only regions with SNPs with modest marginal statistically significance or to screen for any potential false positive credible sets before interpreting mJAM-SuSiE’s credible sets after estimation.

We also carried out a case study of prostate cancer where mJAM is applied to several prostate cancer susceptible regions. Through three different regions with different characteristics in number of estimated independent signals and underlying LD within and between populations, we demonstrated the practical advantages of mJAM-Forward, including allowing more than one causal variant within a region, outputting individual credible sets corresponding to each index, and easily interpretable index variants with conditional estimates. In addition to the three applied examples shown here, mJAM has been applied to perform index variants selection across all regions in the latest multi-population prostate cancer GWAS^37^ which is currently under review.

For all approaches that use marginal summary statistics and reference data, careful consideration and construction of the correlation matrices is important. This includes using a reference panel with ancestry and LD that matches the population in which the original marginal summary statistics were estimated^41,43^. The methods also require that the correlation matrix used is full rank and positive-definite which is often driven by the sample size of the data and the frequency of the SNPs. For mJAM such consideration must be considered across all populations used in the analysis. Firstly, for rare variants, mJAM estimates of multi-population effect and standard errors that can be different from the marginal meta-analyzed estimates which use inverse-variance weighting. mJAM estimation from summary statistics assume Hardy-Weinberg equilibrium which some variants, especially rare variants, might not satisfy. In addition, many rare variants will also have large effect sizes and large standard errors from the population-specific summary statistics thus resulting in more uncertainty in multi-population analysis compared to variants that are common across all populations. Secondly, in regions with extremely significant lead variants from a well-powered GWAS, even small degrees of LD can pull the marginal and conditional effect estimates of other variants away from the null. Thus, false positive signals might be selected if we apply the same threshold for index SNP selection and LD pruning. For such regions, researchers may consider setting a higher significance threshold for secondary signal selection and a more stringent LD threshold for pruning out correlated signals.

In conclusion, mJAM offers a flexible and efficient modeling framework for multi-population fine-mapping that first selects index variants and then constructs credible sets. One key assumption in mJAM is that causal variants and their effect sizes are similar across all populations and there exists evidence suggesting that common causal variants tend to have consistent effect sizes across populations^26–28^. In future research, we plan to relax the current mJAM assumption to allow different true effect sizes across populations. Other potential future directions include follow-up functional analyses based on mJAM credible sets and polygenic risk score models based on mJAM fine-mapped results. mJAM is currently available as a R package for fine-mapping of specific regions and can easily be adapted for genome-wide applications.

## Declaration of interests

The authors declare no competing interests.

## Acknowledgements

This research was supported by P01CA196569, U01CA261339, P30CA014089, U01CA257328, U19CA214253, and R01HG010297.

## Data and code availability

Both mJAM-Forward and mJAM-SuSiE are available as an R package at https://github.com/USCbiostats/hJAM/R. The codes used for simulations and real data examples are available at https://github.com/USCbiostats/hJAM/manuscript_codes.

## Notes

### Competing Interest Statement

The authors have declared no competing interest.

