## Supplemental method and figures for "Fine-Mapping and Credible Set Construction using a Multi-population Joint Analysis of Marginal Summary Statistics from Genome-wide Association Studies"

### Supplemental Material and Methods

#### Multi-ethnic model setup

To simplify notation and without loss of generality, we consider the scenario with three ethnic groups and we are interested in a common set of $p$ SNPs among these ethnic groups. Within each ethnic group, a linear phenotypic model is assumed.

$$\boldsymbol{y}^{(i)} = \boldsymbol{G}^{(i)}\boldsymbol{\beta}_{global} + \boldsymbol{\epsilon}, \text{ for } i = 1,2,3 \quad (1)$$

$$E[\boldsymbol{y}^{(i)}] = \boldsymbol{G}^{(i)}\widehat{\boldsymbol{\beta}}^{(i)}, \text{ for } i = 1,2,3 \quad (2)$$

$$\widehat{\boldsymbol{\beta}}^{(i)} = \boldsymbol{I_p}\boldsymbol{\beta}_{global} + \boldsymbol{\delta}, \text{ for } i = 1,2,3 \quad (3)$$

where

- $N^{(i)}$ is the sample size for the $i^{th}$ ethnic group
- $\boldsymbol{y^{(i)}}$ is a $N^{(i)} \times 1$ vector, denoting the mean-centered phenotypic trait value
- $\boldsymbol{G^{(i)}}$ is a $N^{(i)} \times p$ matrix, the individual-level genotype data where each SNP has been centered to its mean
- $\epsilon \sim \mathrm{N}(0, \sigma^2)$
- $\boldsymbol{\beta_{global}}$ is a $p \times 1$ vector, denoting the conditional effect of the common SNPs. It is assumed that all ethnic groups have the same $\boldsymbol{\beta_{global}}$
- $\widehat{\boldsymbol{\beta}}^{(i)}$ is a $p \times 1$ vector, denoting the conditional effect estimates for the $i^{th}$ ethnic group

Substituting (2) and (3) into (1), we have

$$\begin{pmatrix} \boldsymbol{y}^{(1)} \\ \boldsymbol{y}^{(2)} \\ \boldsymbol{y}^{(3)} \end{pmatrix} = \begin{pmatrix} \boldsymbol{G}^{(1)} & 0 & 0 \\ 0 & \boldsymbol{G}^{(2)} & 0 \\ 0 & 0 & \boldsymbol{G}^{(3)} \end{pmatrix} \begin{pmatrix} \boldsymbol{I_P} \\ \boldsymbol{I_P} \\ \boldsymbol{I_P} \end{pmatrix} \boldsymbol{\beta}_{global} + \boldsymbol{\epsilon}' \quad (4)$$

#### mJAM likelihood with summary statistics

Denote $\boldsymbol{G}_c := \begin{pmatrix} \boldsymbol{G}^{(1)} & 0 & 0 \\ 0 & \boldsymbol{G}^{(2)} & 0 \\ 0 & 0 & \boldsymbol{G}^{(3)} \end{pmatrix}$, $\boldsymbol{y}_c := \begin{pmatrix} \boldsymbol{y}^{(1)} \\ \boldsymbol{y}^{(2)} \\ \boldsymbol{y}^{(3)} \end{pmatrix}$ and $\boldsymbol{I}_c := \begin{pmatrix} \boldsymbol{I_P} \\ \boldsymbol{I_P} \\ \boldsymbol{I_P} \end{pmatrix}$. Using (4) we can derive the distribution of $\boldsymbol{G}_c\boldsymbol{y}_c$ by multiplying the matrix $\boldsymbol{G}_c\boldsymbol{I}_c$ on every term, then we have:

$$\boldsymbol{G}_c\boldsymbol{I}_c\boldsymbol{y}_c \sim MVN\left(((\boldsymbol{G}_c\,\boldsymbol{I}_c)'\boldsymbol{G}_c\,\boldsymbol{I}_c)\boldsymbol{\beta}_{global},\ \sigma^2((\boldsymbol{G}_c\,\boldsymbol{I}_c)'\boldsymbol{G}_c\,\boldsymbol{I}_c)\right) \quad (4)$$

**Fine mapping by mJAM sufficient statistics**

We impose the Zellner's informative g-prior[1] on $\boldsymbol{\beta}_{global}$:

$$\boldsymbol{\beta}_{global}|\sigma^2, \boldsymbol{G}_c, \boldsymbol{y}_c \sim N\left(\frac{g}{g+1}\widehat{\boldsymbol{\beta}}_{global}, \frac{\sigma^2 g}{g+1}((\boldsymbol{G}_c\,\boldsymbol{I}_c)'\boldsymbol{G}_c\,\boldsymbol{I}_c)^{-1}\right)$$

$$\sigma^2|\boldsymbol{G}_c, \boldsymbol{y}_c \sim IG\left(\frac{n}{2}, \frac{s^2}{2} + \frac{1}{2(g+1)}\left(\widehat{\boldsymbol{\beta}}_{global}\right)^T(\boldsymbol{G}_c\,\boldsymbol{I}_c)'\boldsymbol{G}_c\,\boldsymbol{I}_c\,\widehat{\boldsymbol{\beta}}_{global}\right)$$

where $\sigma^2$ is the residual variance in Eq (1), $s^2 = \left(\boldsymbol{y}_c - \boldsymbol{G}_c\,\boldsymbol{I}_c\widehat{\boldsymbol{\beta}}_{global}\right)^T\left(\boldsymbol{y}_c - \boldsymbol{G}_c\,\boldsymbol{I}_c\widehat{\boldsymbol{\beta}}_{global}\right) = (n-k-1)\hat{\sigma}^2$ is the sum of squares, $g$ is a hyperparameter.

Then the marginal posterior distribution of $\boldsymbol{\beta}_{global}$ is:

$$\boldsymbol{\beta}_{global}|\,\boldsymbol{G}_c, \boldsymbol{y}_c \sim T_p\left(n, \frac{g}{g+1}\widehat{\boldsymbol{\beta}}_{global}, \frac{g\left(s^2 + \left(\widehat{\boldsymbol{\beta}}_{global}\right)^T(\boldsymbol{G}_c\,\boldsymbol{I}_c)'\boldsymbol{G}_c\,\boldsymbol{I}_c\,\widehat{\boldsymbol{\beta}}_{global}/(g+1)\right)}{n(g+1)}((\boldsymbol{G}_c\,\boldsymbol{I}_c)'\boldsymbol{G}_c\,\boldsymbol{I}_c)^{-1}\right)$$

And the posterior mean and variance of $\boldsymbol{\beta}_{global}$ is:

$$E[\boldsymbol{\beta}_{global}|\,\boldsymbol{G}_c, \boldsymbol{y}_c] = \frac{g}{g+1}\widehat{\boldsymbol{\beta}}_{global}$$

$$Var[\boldsymbol{\beta}_{global}|\,\boldsymbol{G}_c, \boldsymbol{y}_c] = \frac{g\left(s^2 + \left(\widehat{\boldsymbol{\beta}}_{global}\right)^T(\boldsymbol{G}_c\,\boldsymbol{I}_c)'\boldsymbol{G}_c\,\boldsymbol{I}_c\,\widehat{\boldsymbol{\beta}}_{global}/(g+1)\right)}{n(g+1)}((\boldsymbol{G}_c\,\boldsymbol{I}_c)'\boldsymbol{G}_c\,\boldsymbol{I}_c)^{-1}$$

Following Fernández et al.'s work[2], we recommend setting $g = n$, the total sample size of GWAS, for model simplicity. When the total sample size is relatively large, such choice of $g$ will put more weight on the maximum likelihood estimates of $\boldsymbol{\beta}_{global}$, $\widehat{\boldsymbol{\beta}}_{global}$. As $n \to \infty$, the posterior mean of $\boldsymbol{\beta}_{global}$ will converge to $\widehat{\boldsymbol{\beta}}_{global}$ and the posterior variance of $\boldsymbol{\beta}_{global}$ will converge to $var\left(\widehat{\boldsymbol{\beta}}_{global}\right) = \sigma^2((\boldsymbol{G}_c\,\boldsymbol{I}_c)'\boldsymbol{G}_c\,\boldsymbol{I}_c)^{-1}$.

The first step of mJAM-Forward (Algorithm 1 in main text) is to select index variants based on conditional significance of all variants in a region. This conditional significance is defined by the p-value under the above $g$ prior formulation by calculating a critical value (i.e. estimated effect size divided by its standard error) based on the $g$ prior posterior. By adopting a $g$ prior on $\boldsymbol{\beta}_{global}$, the relative significance of all the variants in the region remain consistent with frequentist approaches but are more robust for estimation as the posterior mean effect $E[\boldsymbol{\beta}_{global}|\,\boldsymbol{G}_c, \boldsymbol{y}_c]$ has a shrinkage factor $\frac{g}{g+1}$ that can prevent issues of overfitting.

**Estimation of sufficient statistics using summary data**

Based on the above expression of the posterior mean and variance of $\boldsymbol{\beta}_{global}$, it is straightforward that the posterior inference on $\boldsymbol{\beta}_{global}$ only depends on $(\boldsymbol{G}_c\,\boldsymbol{I}_c)'\boldsymbol{G}_c\,\boldsymbol{I}_c$, $\boldsymbol{G}_c\boldsymbol{I}_c\boldsymbol{y}_c$, and residual sum of squares $\boldsymbol{y}_c^T\boldsymbol{y}_c$. By expanding the matrices in each term, we can obtain the following summary statistics in the modified mJAM likelihood:

- The $p \times p$ matrix, $(\boldsymbol{G}_c\,\boldsymbol{I}_c)'\boldsymbol{G}_c\,\boldsymbol{I}_c = \boldsymbol{I}_c{}'\boldsymbol{G}_c{}'\boldsymbol{G}_c\boldsymbol{I}_c = (\boldsymbol{G}^{(1)\prime} \quad \boldsymbol{G}^{(2)\prime} \quad \boldsymbol{G}^{(3)\prime})\begin{pmatrix}\boldsymbol{G}^{(1)}\\ \boldsymbol{G}^{(2)}\\ \boldsymbol{G}^{(3)}\end{pmatrix} = \sum_{i=1}^{3}\boldsymbol{G}^{(i)\prime}\boldsymbol{G}^{(i)}$
- The p-vector, $(\boldsymbol{G}_c\,\boldsymbol{I}_c)'\begin{pmatrix}\boldsymbol{y}^{(1)}\\ \boldsymbol{y}^{(2)}\\ \boldsymbol{y}^{(3)}\end{pmatrix} = (\boldsymbol{G}^{(1)\prime} \quad \boldsymbol{G}^{(2)\prime} \quad \boldsymbol{G}^{(3)\prime})\begin{pmatrix}\boldsymbol{y}^{(1)}\\ \boldsymbol{y}^{(2)}\\ \boldsymbol{y}^{(3)}\end{pmatrix} = \sum_{i=1}^{3}\boldsymbol{G}^{(i)\prime}\boldsymbol{y}^{(i)}$
- The sum of squares, $(\boldsymbol{y}^{(1)\prime} \quad \boldsymbol{y}^{(2)\prime} \quad \boldsymbol{y}^{(3)\prime})\begin{pmatrix}\boldsymbol{y}^{(1)}\\ \boldsymbol{y}^{(2)}\\ \boldsymbol{y}^{(3)}\end{pmatrix} = \sum_{i=1}^{3}\boldsymbol{y}^{(i)\prime}\boldsymbol{y}^{(i)}$
- The sample size of each GWAS, $N^{(1)}, N^{(2)}, N^{(3)}$

Thus, we can estimate $\boldsymbol{G}^{(i)\prime}\boldsymbol{G}^{(i)}$, $\boldsymbol{G}^{(i)\prime}\boldsymbol{y}^{(i)}$, $\boldsymbol{y}^{(i)\prime}\boldsymbol{y}^{(i)}$ for each population individually and then sum up the population-specific statistics to obtain the mJAM summary statistics. To simply notions, we will drop the superscripts and just use $\boldsymbol{G}'\boldsymbol{G}, \boldsymbol{G}'\boldsymbol{y}, \boldsymbol{y}'\boldsymbol{y}$ in the following discussion.

**Estimation of $\boldsymbol{G}'\boldsymbol{G}$**

Following Yang *et al.*'s work[3], the variance-covariance matrix of the GWAS samples, $\boldsymbol{G}'\boldsymbol{G}$, can be estimated from the LD structure from a reference panel, and effect allele frequencies from the GWAS summary statistics data.

Let $\boldsymbol{W} = \{w_{ij}\}$ be the genotype matrix of the reference panel with sample size m. Standardize $W$ by letting $w_{ij} = -2f_i, 1-2f_i, 2-2f_i$, where $f_i$ is the allele frequency of SNP $i$ in the reference data. Due to the difference in sample sizes of the reference data and the GWAS data, we cannot use $\boldsymbol{W}'\boldsymbol{W}$ to estimate $\boldsymbol{G}'\boldsymbol{G}$ directly. To adjust for such difference, firstly define

$$\boldsymbol{B} = \boldsymbol{D}^{1/2}\boldsymbol{D}_{\boldsymbol{w}}^{-1/2}\boldsymbol{W}'\,\boldsymbol{W}\,\boldsymbol{D}_{\boldsymbol{w}}^{-1/2}\,\boldsymbol{D}^{1/2} \qquad (5)$$

where $\boldsymbol{D} = \{D_j\}$ is the diagonal matrix of $\boldsymbol{G}'\boldsymbol{G}$, with $D_j = \sum_{j=1}^{N^{(i)}} g_{ij}^2$. And $\boldsymbol{D}_{\boldsymbol{W}}$ is the diagonal matrix of $\boldsymbol{W}'\boldsymbol{W}$, with $D_{W(j)} = \sum_{j=1}^{m} w_{ij}^2$. $\boldsymbol{B}$ is then an estimate for the variance-covariance matrix of the GWAS samples $\boldsymbol{G}'\boldsymbol{G}$. Since $D_j = \sum_{j=1}^{N^{(i)}} g_{ij}^2$ are still unknown, we use $D_j = 2p_j(1-p_j)N$ to estimate $\sum_{j=1}^{N^{(i)}} g_{ij}^2$ under HWE, where $p_j$ is the allele frequency of SNP $j$ in the GWAS sample.

### Estimation of $\boldsymbol{G'y}$

The total trait burden of each SNP, $\boldsymbol{G'y}$, can be estimated from effect allele frequencies and marginal effect estimates when HWE is assumed.[2] Define $\boldsymbol{z} := \boldsymbol{G'y}$. Let $\hat{n}_{j0}, \hat{n}_{j1}, \hat{n}_{j2}$ denote the group counts of genotype = 0,1,2 for SNP $j$ respectively. For SNP $j$,

$$z_j = \bar{y}_{j1} n_{j1} + 2\bar{y}_{j2} n_{j2} \qquad (6)$$

Assuming Hardy-Weinberg equilibrium (HWE), we have $\hat{n}_{j0} = \left(1 - \hat{p}_j\right)^2 N$, $\hat{n}_{j1} = 2\,\hat{p}_j\left(1 - \hat{p}_j\right)N$, and $\hat{n}_{j2} = \hat{p}_j^2 N$, where $\hat{p}_j$ denotes the allele frequency of SNP $j$ and $N$ is the sample size of the GWAS summary data. And $\bar{y}_{j1}$, $\bar{y}_{j2}$ can be estimated under a linear phenotypic model as

$\hat{\bar{y}}_{j0} = -\frac{\hat{n}_{j1}\hat{\beta}_j + 2\hat{n}_{j2}\hat{\beta}_j}{N}$, $\hat{\bar{y}}_{j1} = \hat{\bar{y}}_{j0} + \hat{\beta}_j$, and $\hat{\bar{y}}_{j2} = \hat{\bar{y}}_{j0} + 2\hat{\beta}_j$. Substituting all estimates into (6), we get $\boldsymbol{z} = \boldsymbol{G'y}$.

### Estimation of $\boldsymbol{y'y}$

In the association analysis of a single SNP $j$, the estimate of $\boldsymbol{y'y}$ is provided by

$$\boldsymbol{y'y} = D_j S_j^2\left(N^{(i)} - 1\right) + D_j \hat{\beta}_j^2 \qquad (7)$$

where $\hat{\beta}_j$ is the marginal effect estimate for SNP $j$ and $S_j^2$ is the squared standard error of $\hat{\beta}_j$.[1] Yang *et al.* suggested to take the median of $D_j S_j^2\left(N^{(i)} - 1\right) + D_j \hat{\beta}_j^2$ across all SNPs to get a single estimate of $\boldsymbol{y'y}$ for the $i^{th}$ ethnic group. However, for rare SNPs whose estimates that have relatively large standard errors, the median estimate of $\boldsymbol{y'y}$ tends to underestimate the residual variance in the one-SNP models. Thus, for all one-SNP models in mJAM, we propose a modified estimate of $\boldsymbol{y'y}$ that takes a weighted average of median and individual estimates.

$$\boldsymbol{y'y} = w_j \cdot \boldsymbol{y'y}_{\boldsymbol{j}} + (1 - w_j) \cdot \boldsymbol{y'y}_{\boldsymbol{m}} \qquad (8)$$

where $\boldsymbol{y'y}_{\boldsymbol{j}} = \boldsymbol{y'y} = D_j S_j^2\left(N^{(i)} - 1\right) + D_j \hat{\beta}_j^2$ for SNP $j$, $\boldsymbol{y'y}_{\boldsymbol{m}}$ is the median across all $\boldsymbol{y'y}_{\boldsymbol{j}}$, $w_j = \frac{\boldsymbol{y'y}_{\boldsymbol{m}}}{\boldsymbol{y'y}_{\boldsymbol{m}} + |\boldsymbol{y'y}_{\boldsymbol{m}} - \boldsymbol{y'y}_{\boldsymbol{j}}|}$ is the relative weight of median $\boldsymbol{y'y}$ to SNP-specific $\boldsymbol{y'y}$.

### Posterior Model Probability

To quantify how significant is a putative credible set SNP associated with the outcome, $\boldsymbol{Y}$, we adopted a Bayesian posterior probability of models built upon Bayes factors for pairs of hypotheses.[3] For a putative credible set SNP denoted as $j$, the posterior probability of the one-SNP model of $j, M_j$, can be expressed as

$$Pr\left(M_j \mid Data\right) = \frac{p(M_j)BF\left[M_j: M_{Null}\right]}{\sum_j p(M_j)BF\left[M_j: M_{Null}\right]}$$

where $p(M_j)$ is the prior probability of model $M_j$, and $BF\left[M_j: M_{Null}\right]$ is the Bayes factor for comparing $M_j$ to $M_{Null}$, the null model. $BF\left[M_j: M_{Null}\right]$ is simply the ratio of marginal likelihood of the data under model $M_j$ versus that under the null model:

$$BF\left[M_j: M_{Null}\right] = \frac{P(\boldsymbol{Y}|M_j)}{P(\boldsymbol{Y}|M_{Null})}$$

We further adopt Zellner's g prior in the derivation of $BF\left[M_j: M_{Null}\right]$. Zellner's g prior is a conjugate Normal-Gamma prior for linear regression models $\boldsymbol{Y} = \boldsymbol{G\beta} + \epsilon$ where

$$\boldsymbol{\beta}|\phi \sim N\left(\widetilde{\boldsymbol{\beta}}, g\sigma^2(\boldsymbol{G}'\boldsymbol{G})^{-1}\right), p(\sigma^2) \propto \sigma^{-2}$$

and $\widetilde{\boldsymbol{\beta}}$ is the prior mean effect which is usually set to 0 in genetic models.

It has shown that the Bayes factor under g prior formulation comparing $M_j$ to the null model, $BF\left[M_j: M_{Null}\right]$, can be written as

$$BF\left[M_j: M_{Null}\right] = \left[\frac{1+g}{1+g(1-R_j^2)}\right]^{(n-1)/2} (1+g)^{-p_j/2}$$

where $R_j^2$ is the coefficient of determination of regression model $M_j$, $n$ is the sample size in $M_j$, and $p_j$ is the number of coefficients in $M_j$. [3]

With only summary statistics, the coefficient of determination $R_j^2$ can be expressed as

$$R_j^2 = \frac{\widehat{\boldsymbol{b}}'\boldsymbol{G}'\boldsymbol{y}}{\boldsymbol{y}'\boldsymbol{y}} = \frac{(\boldsymbol{G}'\boldsymbol{y})'(\boldsymbol{G}'\boldsymbol{G})^{-1}\boldsymbol{G}'\boldsymbol{y}}{\boldsymbol{y}'\boldsymbol{y}}$$
$$= \frac{\widehat{\boldsymbol{b}}'\boldsymbol{D}\widehat{\boldsymbol{\beta}}}{\boldsymbol{y}'\boldsymbol{y}} = \frac{(\boldsymbol{D}\widehat{\boldsymbol{\beta}})'(\boldsymbol{G}'\boldsymbol{G})^{-1}\boldsymbol{D}\widehat{\boldsymbol{\beta}}}{\boldsymbol{y}'\boldsymbol{y}}$$

where $\boldsymbol{G}'\boldsymbol{G}$ is variance-covariance matrix of SNPs within $M_j$, $\boldsymbol{D}$ is the diagonal matrix of $\boldsymbol{G}'\boldsymbol{G}$**,** and $\widehat{\boldsymbol{\beta}}$ is the vector of marginal effect estimates of SNPs within $M_j$. All these components can be estimated using the marginal summary statistics with the approach discussed in previous sections.

To build the credible sets for index SNPs selected conditional on the first index SNP in the region, we adjust the posterior model probabilities of putative credible set SNPs to also be conditional on the presence of any previous index SNP(s). Then for a putative credible set SNP $j$, model $M_j$ is now a multi-SNP model with SNP $j$ and all previous index SNP(s). We then replace $R_j^2$ with partial $R^2$ of SNP $j$ conditional on all previous index SNP(s) to calculate $BF[M_j\colon M_{Null}]$, that is,

$$R^2_{\gamma_2|\gamma_1} = \frac{SSR(\gamma_2) - SSR(\gamma_1)}{SSTO - SSR(\gamma_1)} = \frac{(\widehat{\boldsymbol{b}}'\boldsymbol{D}\widehat{\boldsymbol{\beta}})_{\gamma_2} - (\widehat{\boldsymbol{b}}'\boldsymbol{D}\widehat{\boldsymbol{\beta}})_{\gamma_1}}{\boldsymbol{y}'\boldsymbol{y} - (\widehat{\boldsymbol{b}}'\boldsymbol{D}\widehat{\boldsymbol{\beta}})_{\gamma_1}}$$

where $\gamma_1$ is the model with all previous index SNP(s), $\gamma_2$ is the model with SNP $j$ and all previous index SNP(s). Note that $\gamma_1$ is nested within $\gamma_2$ and $\widehat{\boldsymbol{b}}$ refers to the joint effect estimates which can be obtained using the approach discussed in previous sections.

**Posterior Mediation Probability**

As shown in Figure 1 in the main text, the mediation effect of an index SNP, X, on the relationship between a candidate credible set SNP, W, and the outcome, Y, can be evaluated through the difference in the total effect and the indirect effect, i.e. $|\tau_W - \tau'_W|$, where $\tau_W$ is the total effect (the marginal effect of W on Y) and $\tau'_W$ is the indirect effect (the adjusted effect of W on Y adjusted for X). Then the mediation probability is expressed as

$$\Pr(Mediation|Data) = \Pr(|\tau_W - \tau'_W| > 0|Data)$$

We again adopt Zellner's g prior formulation and obtain a Wald-type statistics for $|\tau_W - \tau'_W|$. Since the g prior is a conjugate prior, we can express the posterior distribution of $\boldsymbol{\beta}$ and $\sigma^2$ in closed forms, that is

$$\boldsymbol{\beta}|\sigma^2, \boldsymbol{y}, \boldsymbol{G} \sim N\left(\frac{g}{g+1}\left(\frac{\widetilde{\boldsymbol{\beta}}}{g} + \widehat{\boldsymbol{\beta}}\right), \frac{\sigma^2 g}{g+1}(\boldsymbol{G}^T\boldsymbol{G})^{-1}\right)$$

$$\sigma^2|\boldsymbol{y}, \boldsymbol{G} \sim IG\left(\frac{n}{2}, \frac{s^2}{2} + \frac{1}{2(g+1)}\left(\widetilde{\boldsymbol{\beta}} - \widehat{\boldsymbol{\beta}}\right)^T \boldsymbol{G}^T\boldsymbol{G}\left(\widetilde{\boldsymbol{\beta}} - \widehat{\boldsymbol{\beta}}\right)\right)$$

where n is the sample size, $s^2 = \left(\boldsymbol{y} - \boldsymbol{G}\widehat{\boldsymbol{\beta}}\right)^{\boldsymbol{T}}(\boldsymbol{y} - \boldsymbol{G}\widehat{\boldsymbol{\beta}})$, $\widetilde{\boldsymbol{\beta}} = \boldsymbol{0}$ is the prior mean effect, $\widehat{\boldsymbol{\beta}} = (\boldsymbol{G^T G})^{-\boldsymbol{1}}\boldsymbol{G^T y}$ is the maximum likelihood estimator of $\boldsymbol{\beta}$. Then the posterior distribution of $\boldsymbol{\beta}$ without conditioning on $\sigma^2$ is

$$\boldsymbol{\beta} |\, \boldsymbol{y}, \boldsymbol{G} \sim T_p\left(n, \frac{g}{g+1}\left(\frac{\widetilde{\boldsymbol{\beta}}}{g}+\widehat{\boldsymbol{\beta}}\right), \frac{g\left(s^2+\left(\widetilde{\boldsymbol{\beta}}-\widehat{\boldsymbol{\beta}}\right)^T \boldsymbol{G}^T \boldsymbol{G}\left(\widetilde{\boldsymbol{\beta}}-\widehat{\boldsymbol{\beta}}\right)/(g+1)\right)}{n(g+1)}\left(\boldsymbol{G}^T \boldsymbol{G}\right)^{-1}\right)$$

where $p$ is the length of $\boldsymbol{\beta}$ and $T_p$ is the multivariate t distribution of $p$ dimensions and $n$ degrees of freedom. As $n \to \infty$, $T_p$ converges to a multivariate Gaussian distribution of $p$ dimensions.

Assuming the posterior distributions of $\tau_W$ and $\tau_W'$ are independent, and denote the posterior mean of $\tau_W$ is $\tau_a$ and the variance in the posterior distribution of $\tau$ is $v(\tau_a)$. Similarly for $\tau_W'$ , the posterior mean is $\tau_a'$ and the variance is $v(\tau_a')$. Then the Wald-type statistic for $|\tau_W - \tau_W'|$ is

$$Z_\tau \coloneqq \frac{|\tau_a - \tau_a'|}{\sqrt{v(\tau_a)+v(\tau_a')}}$$

where $\tau_a$, $\tau_a'$, $v(\tau_a)$, and $v(\tau_a')$ estimated via the closed form approximate multi-variate Gaussian distribution shown before. Then we evaluate the posterior mediation probability as

$$\Pr(|\tau_W - \tau_W'| > 0 | Data) = \Pr(Z_\tau > 0)$$

where $Z_\tau$ approximately follows a standard normal distribution.

**Multi-ethnic Fine Mapping Using Sum of Single Effect Model (mJAM-SuSiE)**

To allow for multiple non-zero effects in the common joint effect $\boldsymbol{\beta}_{\text{global}}$, the Sum of Single Effect Model (SuSiE) proposed a new approach to model the sparse vector of $\boldsymbol{\beta}_{\text{global}}$ as a sum of "single-effect" vectors, each with one non-zero effect. Taken together the multi-ethnic joint analysis setting (equation (3) in main text) and SuSiE, fine-mapping in terms of the common joint effect $\boldsymbol{\beta}_{\text{global}}$ can be expressed as

$$\boldsymbol{\beta}_{\text{global}} = \sum_{l=1}^{L} \boldsymbol{\beta}_{\boldsymbol{l}} = \sum_{l=1}^{L} \beta_l \boldsymbol{\gamma}_{\boldsymbol{l}} \tag{4}$$

$$\boldsymbol{\gamma}_{\boldsymbol{l}} \sim \text{Mult}(1, \tau) \text{ and } \beta_l \sim N_1\left(0, \sigma_{0l}^2\right), \tag{5}$$

where $l$ is the index of credible sets, and $L$ denotes the largest number of credible sets allowed in fitting, and $\sigma_{0l}^2$ denotes the prior variance of the non-zero effect $\beta_l$. SuSiE inference is robust to overstating $L$; thus, in practice we recommend setting $L$ larger than the number of potential causal signals in a region.

The susieR package [4] provides the implementation of SuSiE not only with inputs as individual-level data, but also with inputs as summary statistics data. When only summary data is available, we employ $(\boldsymbol{G}_c\,\boldsymbol{I}_c)'\boldsymbol{G}_c\,\boldsymbol{I}_c$, $(\boldsymbol{G}_c\,\boldsymbol{I}_c)'\boldsymbol{y}_c$, and $\boldsymbol{y}_c{}'\boldsymbol{y}_c$, which can be estimated from ethnic-specific marginal effect estimates and reference individual-level dosage data for each ethnic group, to obtain the posterior mean and posterior inclusion probability (PIP) of each SNP using the summary statistic version of SuSiE. See Supplimental Supplemental Algorithm 1 below for fitting mJAM-SuSiE model.

**Supplemental Algorithm 1 Pseudo algorithm for fitting mJAM-SuSiE and constructing credible sets using SuSiE PIP.**

Input data: $\widehat{\boldsymbol{\beta}}^{(i)}$, $se\left(\widehat{\boldsymbol{\beta}}^{(i)}\right)$, $N_{GWAS}$, $\mathbf{EAF}^{(\mathbf{i})}$, $\boldsymbol{G}_{\boldsymbol{R}}^{(\boldsymbol{i})}$ for each study indexed by $i$
Input arguments: $L$, the maximum number of credible sets allowed; $r$, the minimum absolute correlation allowed in a credible set; requested coverage.
Function required:
(1) susie_suff_stat$(\boldsymbol{X}'\boldsymbol{X},\ \boldsymbol{X}'\boldsymbol{y};\ L) \rightarrow \boldsymbol{PIP}$ that computes a $p$ by $L$ posterior probability matrix using SuSiE with summary statistics.

1. Compute mJAM statistics $(\boldsymbol{G}_c\,\boldsymbol{I}_c)'\boldsymbol{G}_c\,\boldsymbol{I}_c$, $(\boldsymbol{G}_c\,\boldsymbol{I}_c)'\boldsymbol{y}_c$, and $\boldsymbol{y}_c{}'\boldsymbol{y}_c$
2. Fit susie_suff_stat$(\boldsymbol{X}'\boldsymbol{X},\ \boldsymbol{X}'\boldsymbol{y};\ L) \rightarrow \boldsymbol{PIP}$
3. For $l$ in 1,…, $L$ do
4. Find a set of SNPs where cumulative $\boldsymbol{PIP}^{(l)}$ reaches the requested coverage and all pairwise absolute correlation are no less than $r$. If not, report no credible set.

Return credible set(s) with index SNP(s) and PIP.

**Incorporating missing variants in mJAM**

Suppose SNP $m$ is missing in population $k$. To analyze all available data without filtering this SNP, we can find the identity matrix of population $k$ in the mJAM model, then replace the diagonal term of the corresponding SNP with 0. Now the identify matrix is as:

$$\begin{pmatrix} \boldsymbol{I}_p^{(1)} & 0 & 0 & 0 & 0 \\ 0 & \dots & 0 & 0 & 0 \\ 0 & 0 & \boldsymbol{I}_p^{(k)} & 0 & 0 \\ 0 & 0 & 0 & \dots & 0 \\ 0 & 0 & 0 & 0 & \boldsymbol{I}_p^{(K)} \end{pmatrix}$$

and $\boldsymbol{I}_p^{(k)} = \begin{pmatrix} 1 & & & & & \\ & 1 & & & & \\ & & \ddots & & & \\ & & & 0 & & \\ & & & & \ddots & \\ & & & & & 1 \end{pmatrix}$ where the $m^{th}$ term on the diagonal is replaced by 0.

For SNP $m$, the resulting mJAM statistics become the following:

$$\sum_{i=1}^{K} \boldsymbol{G}_m^{(i)'} \boldsymbol{y}_m^{(i)} = \boldsymbol{G}_m^{(1)'} \boldsymbol{y}_m^{(1)} + \cdots + \boldsymbol{G}_m^{(k-1)'} \boldsymbol{y}_m^{(k-1)} + \boldsymbol{G}_m^{(k+1)'} \boldsymbol{y}_m^{(k+1)} + \cdots + \boldsymbol{G}_m^{(K)'} \boldsymbol{y}_m^{(K)}$$

$$\sum_{i=1}^{K} \boldsymbol{G}_m^{(i)'} \boldsymbol{G}_m^{(i)} = \boldsymbol{G}_m^{(1)'} \boldsymbol{G}_m^{(1)} + \cdots + \boldsymbol{G}_m^{(k-1)'} \boldsymbol{G}_m^{(k-1)} + \boldsymbol{G}_m^{(k+1)'} \boldsymbol{G}_m^{(k+1)} + \cdots + \boldsymbol{G}_m^{(K)'} \boldsymbol{G}_m^{(K)}$$

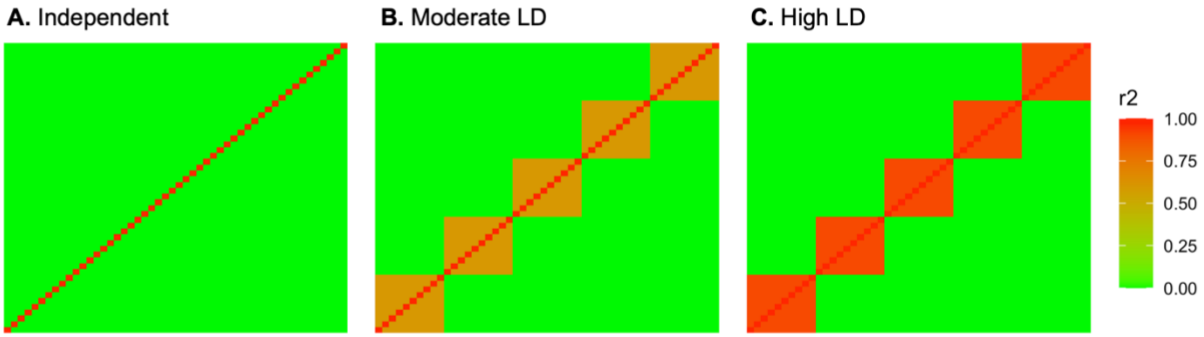


**Figure S 1 LD structure in simulation studies**

*(A) Independent: all SNPs are independent with each other; (B) Moderate LD: 5 blocks of 10 SNPs each where pairwise* $r^2$ *is uniformly set to 0.6; (C) High LD: 5 blocks of 10 SNPs each where pairwise* $r^2$ *is uniformly set to 0.9.*

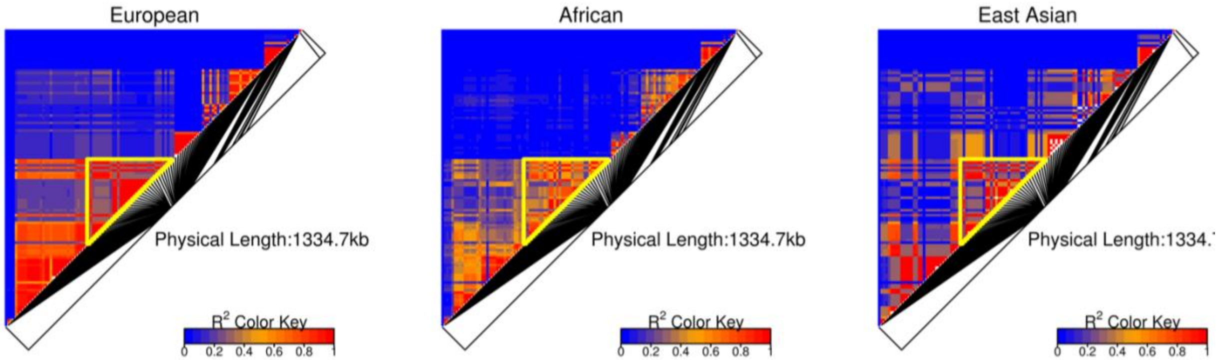


**Figure S 2 LD structure of European, African and East Asian populations in chromosome 2 region from position 168212955 to 169812955.**

*Candidate causal SNP was drawn randomly from the block highlighted in yellow and satisfied meta P-value < 1e-9.*

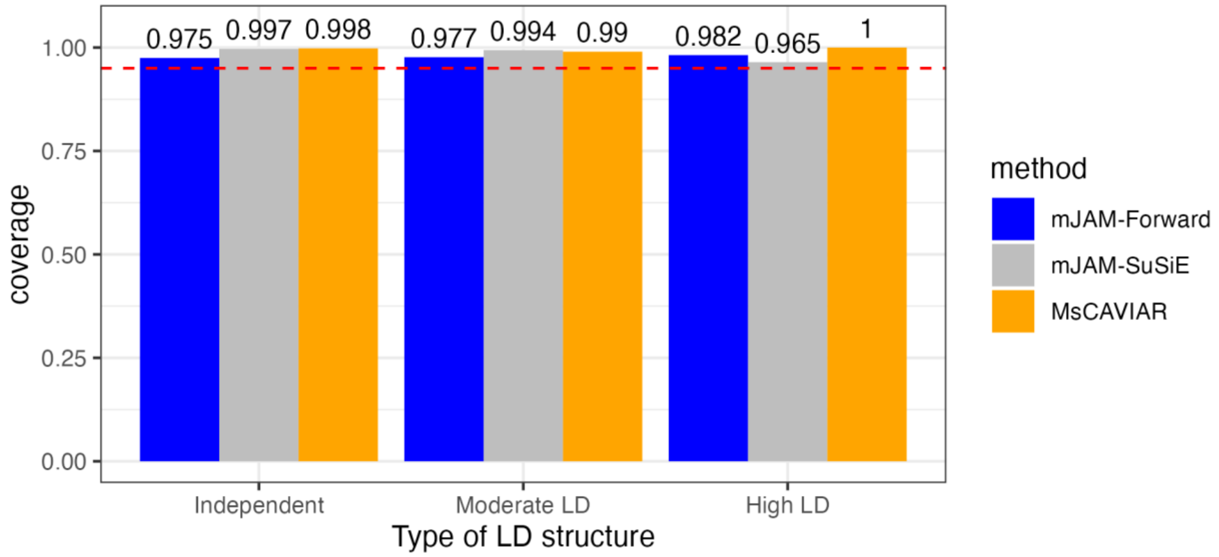


**Figure S 3 Empirical coverage for mJAM-Forward, mJAM-SuSiE and MsCAVIAR 95% credible sets under baseline scenario.**

*Baseline scenario: 50 SNPs in total, 1 causal SNP with an effect size of 0.03, 3 studies per ancestry group, and balanced sample size across populations. For independent LD structure, there is no correlation between SNPs within each LD block. For moderate LD structure, the pairwise r2 between SNPs within each LD block is set to be 0.62 for all populations. For high LD structure, the pairwise r2 between SNPs within each LD block is set to be 0.92 for all populations. Empirical coverage is defined as the observed proportion of 95% credible sets that included at least one true causal SNP. The requested coverage level of 95% is indicated in red dashed lines.*

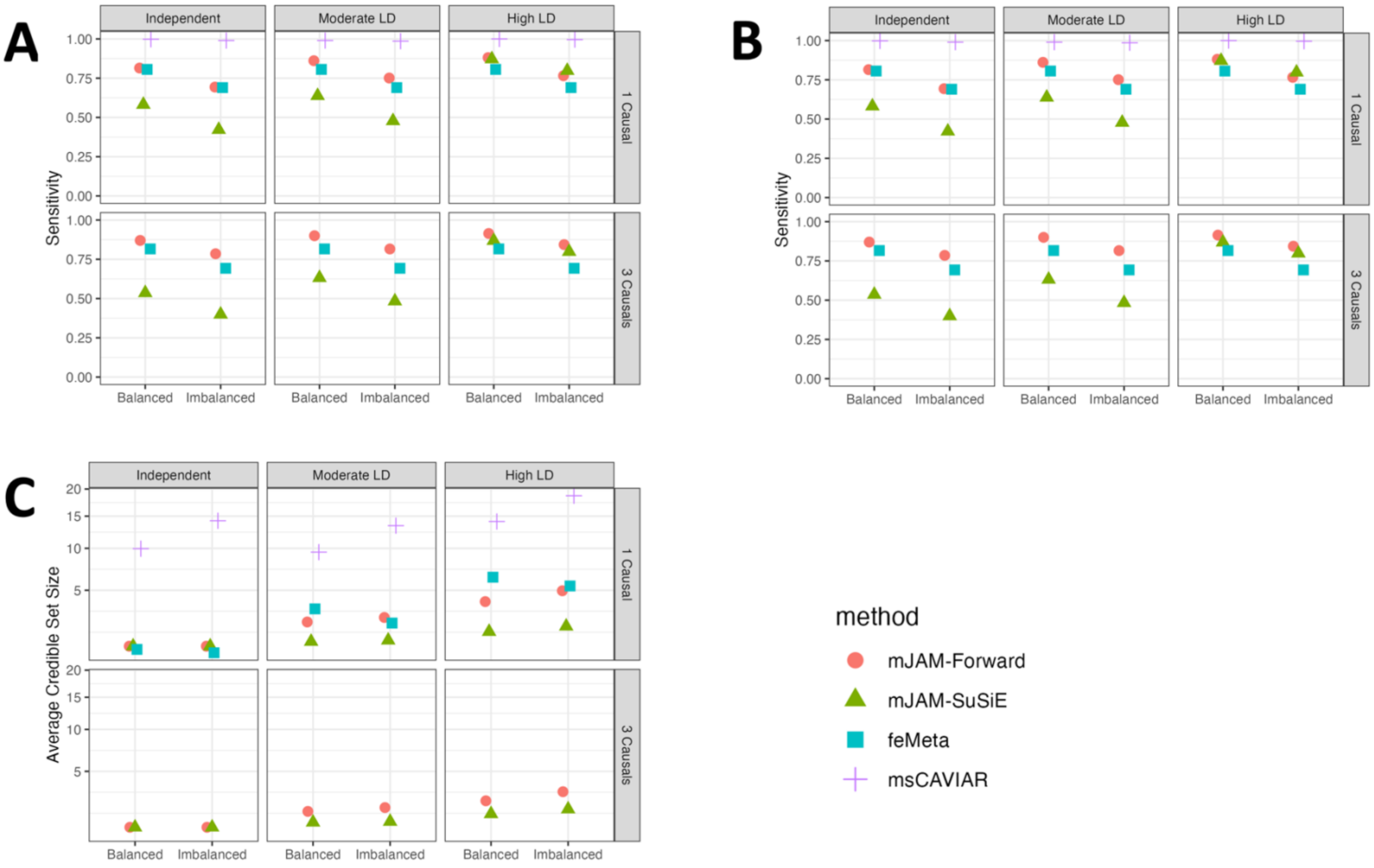


**Figure S 4 Credible set performance in simulation studies with artificial LD structure.**

*For balanced sample size, all 3 populations have a total sample size of 15,000. For unbalanced sample size, the first population has a total sample size of 11,000 and the other two populations have a total sample size of 2,000. Both sample size scenarios have sample size added up to 45,000. For both sample size scenarios, there is either 1 causal SNP or 3 causal SNPs in separate LD blocks, each with an effect size of 0.03; the number of studies per population is set to be 3. (A) Sensitivity, the proportion of true causal SNPs being selected in a credible set, averaged over 500 simulations. (B) PPV, the proportion of true causal SNPs over the total number of selected credible set SNPs, averaged over 500 simulations. (C) Average CS size, the number of SNPs in each 95% credible sets, averaged over 500 simulations.*

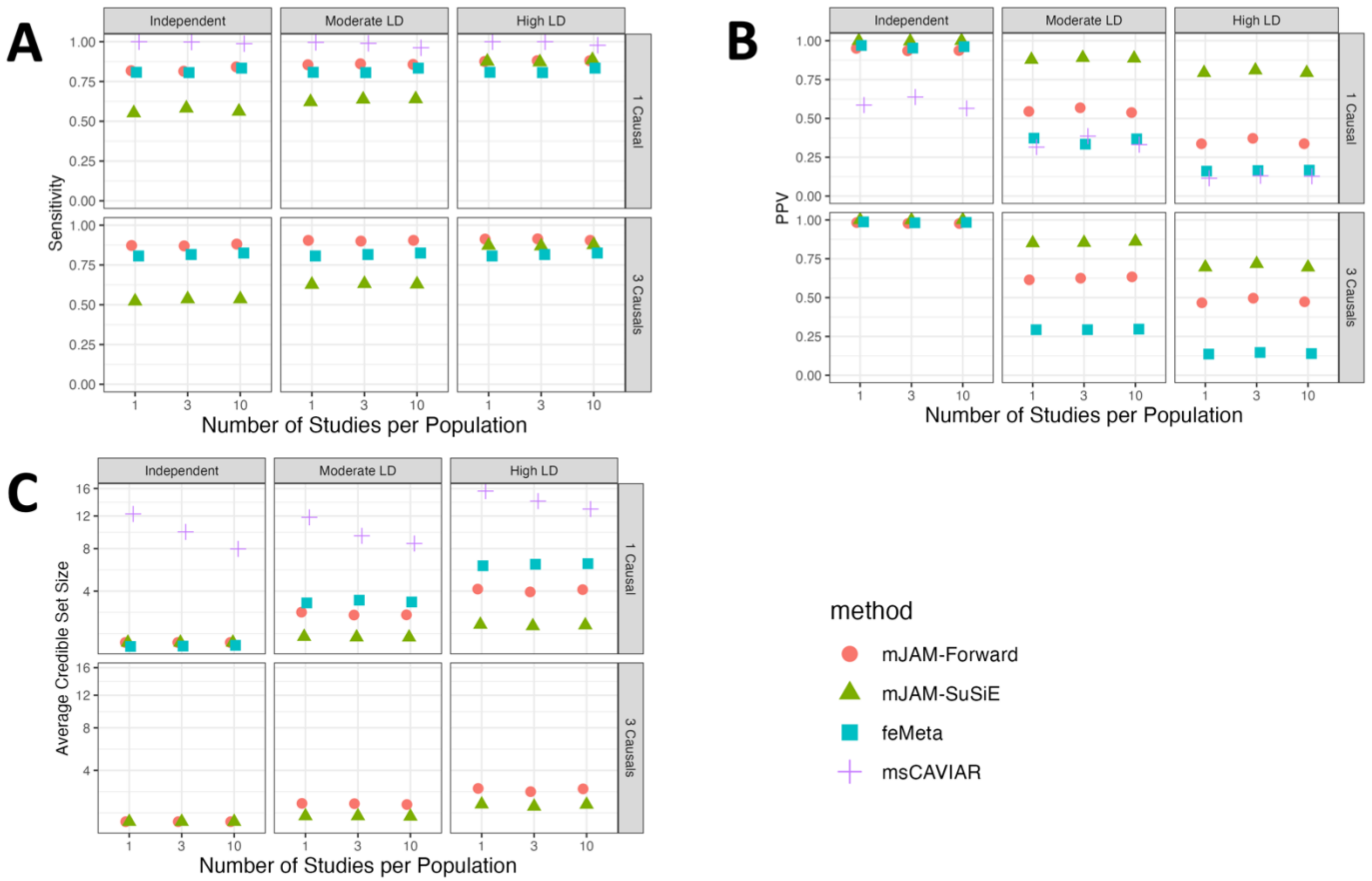


**Figure S 5 Credible set performance in simulation studies with various number of studies from each ancestry group.**

*There are 3 ancestry groups in total, each with 1 or 3 or 10 studies. The total sample size for each ancestry group is fixed at 15,000 and sample size is the same across individual studies within each ancestry group. There is either 1 causal SNP or 3 causal SNPs in separate LD blocks, each with an effect size of 0.03. (A) Sensitivity, the proportion of true causal SNPs being selected in a credible set, averaged over 500 simulations. (B) PPV, the proportion of true causal SNPs over the total number of selected credible set SNPs, averaged over 500 simulations. (C) Average CS size, the number of SNPs in each 95% credible sets, averaged over 500 simulations.*

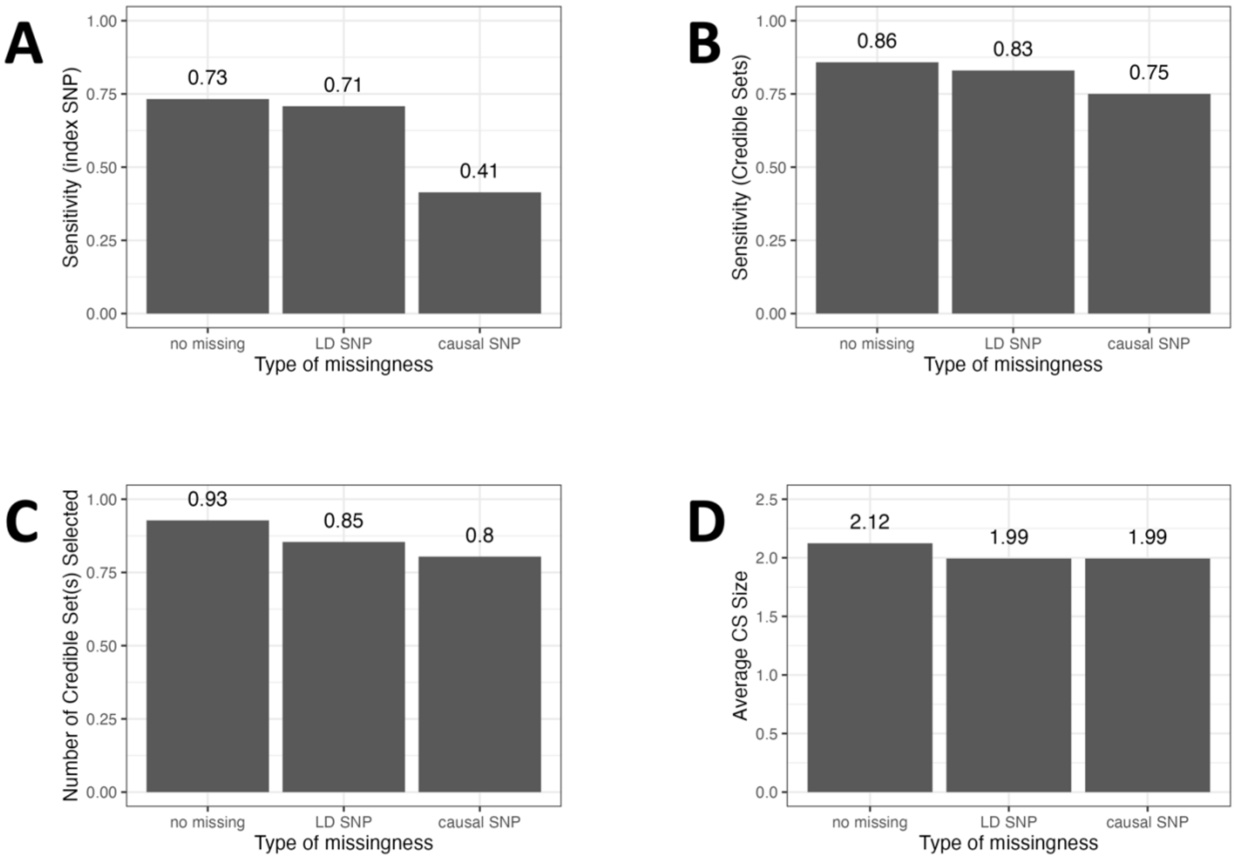


**Figure S 6 Performance of mJAM-Forward in simulation studies with various missingness under baseline scenario.**

*Baseline scenario: 1 causal SNP out of 50 SNPs in total; 3 ancestry groups and 3 studies per ancestry group; sample size = 5,000 per study; pairwise r2 between SNPs within each LD block is 0.60. There are 3 types of missingness simulated: (1) no missing: all SNPs are available in each study; (2) LD SNP missing: one SNP that is in the same LD block with the causal SNP is missing in one study.*

*Note that the other two studies of the same ancestry group have this SNP available. (3) causal SNP: the causal SNP is missing in one study but that the other two studies of the same ancestry group have the causal SNP available. (A) Index SNP sensitivity, the proportion of true causal SNPs being selected as an index SNP, averaged over 500 simulations. (B) Credible set sensitivity, the proportion of true causal SNPs being selected in a credible set, averaged over 500 simulations. (C) Number of 95% credible set(s) selected, averaged over 500 simulations. (D) Average CS size, the number of SNPs in each 95% credible sets, averaged over 500 simulations.*

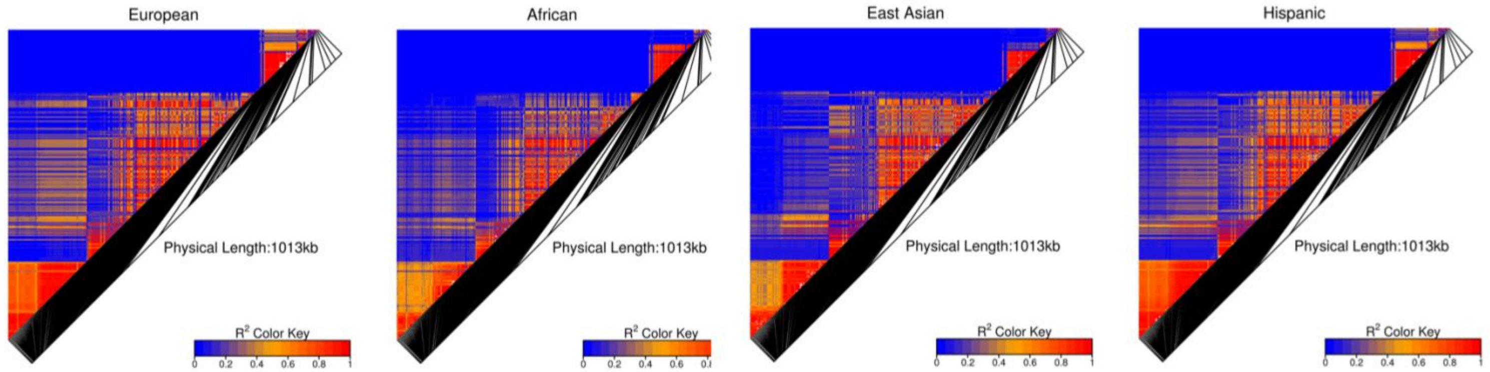


**Figure S 7 Ethnic-specific LD structure for 416 SNPs at chromosome 12 from position 109194870 to 110794870.**

*Analysis restricted to SNPs with meta-analyzed P-value < 0.001 and MAF > 2%.*

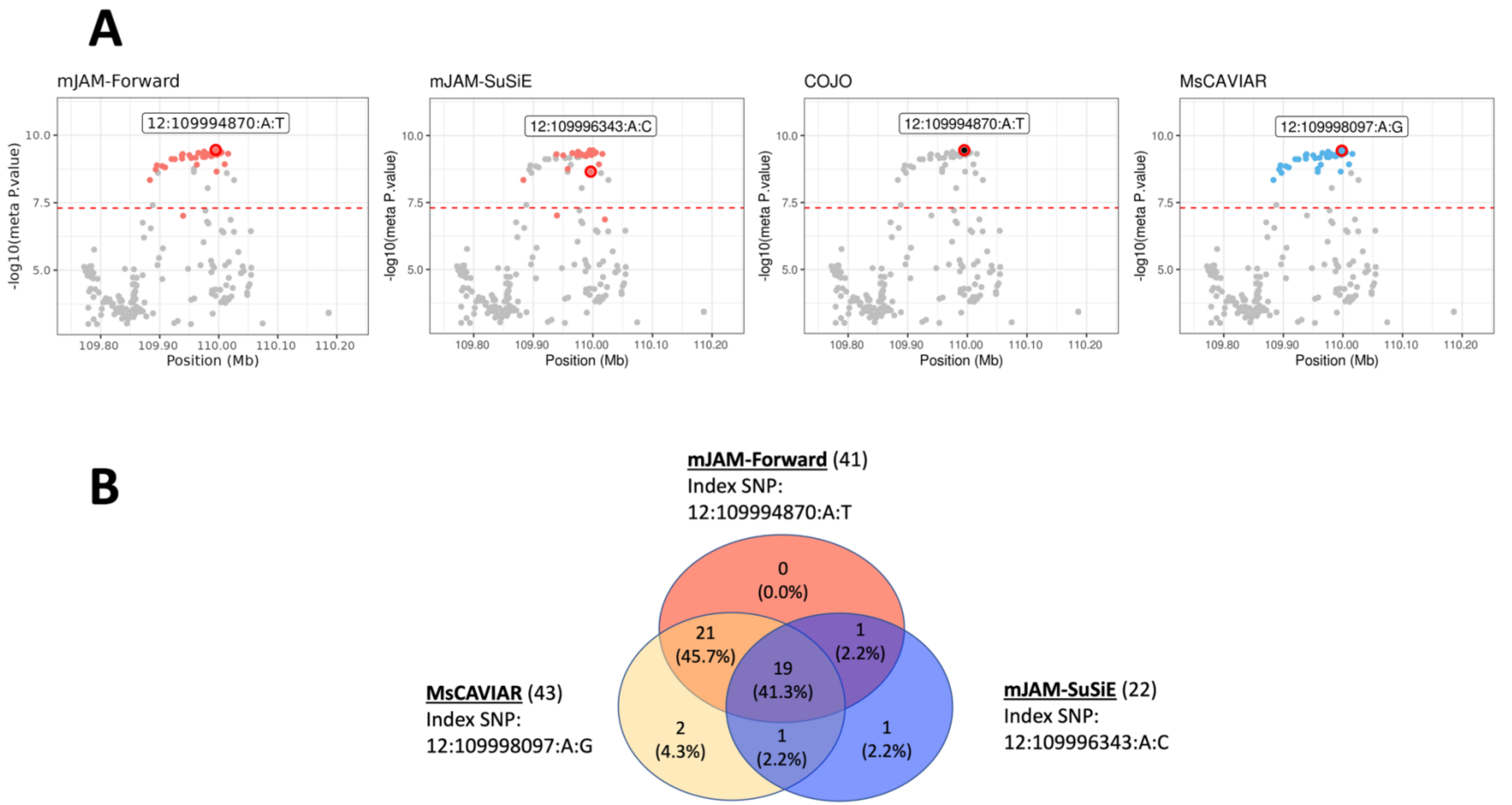


**Figure S 8 Fine-mapping results for chromosome 12 region from position 109194870 to 110794870.**

*(A) From left to right is mJAM-Forward 95% credible set, mJAM-SuSiE 95% credible set, index SNP selected by COJO, and MsCAVIAR 95% credible set. Index SNPs are circled in red. SNPs in the same credible sets are highlighted in the same color. Genome-wide significance (* $5 \times 10^{-8}$ *) is shown in red dashed line. (B) Venn diagram showing the overlap between 95% credible sets from mJAM-Forward, mJAM-SuSiE and MsCAVIAR.*

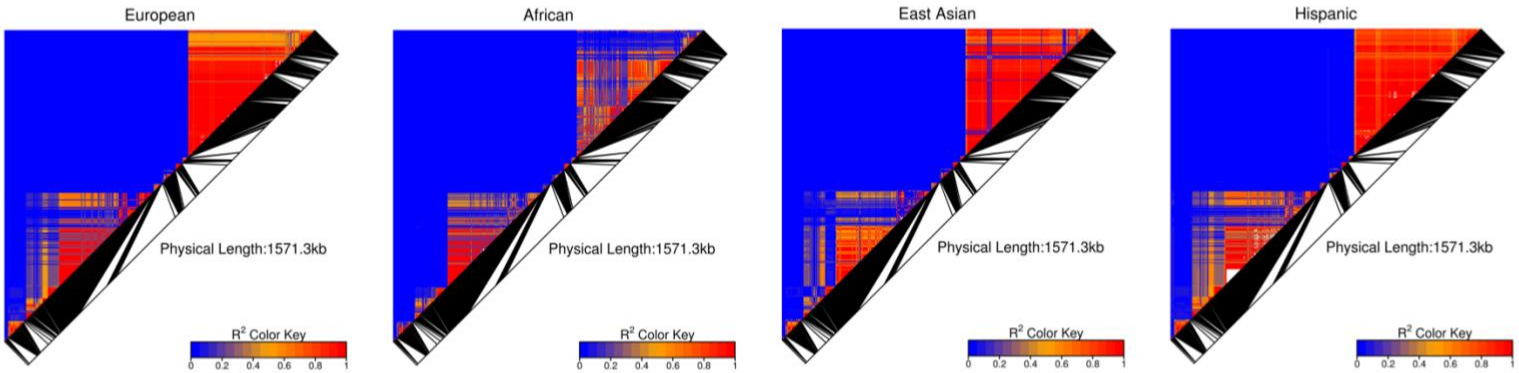


**Figure S 9 Ethnic-specific LD structure for 416 SNPs at chromosome 10 from position 79436999 to 81635998.**

*Analysis restricted to SNPs with meta-analyzed P-value < 0.001 and MAF > 2%.*

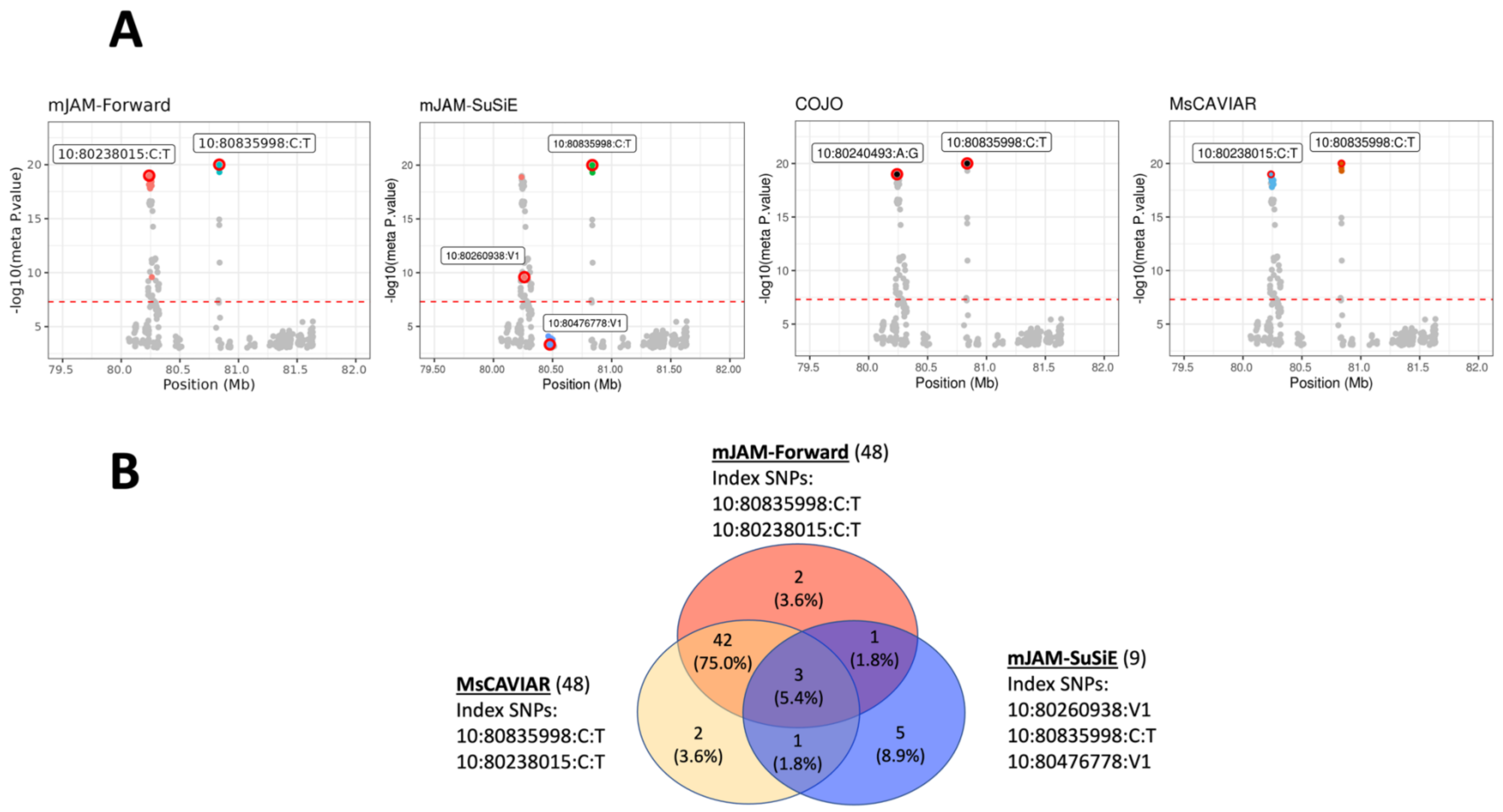


**Figure S 10 Fine-mapping results for chromosome 10 region from position 79436999 to 81635998.**

*(A) From left to right is mJAM-Forward 95% credible set, mJAM-SuSiE 95% credible set, index SNP selected by COJO, and MsCAVIAR 95% credible set. Index SNPs are circled in red. SNPs in the same credible sets are highlighted in the same color. Genome-wide significance ( $5 \times 10^{-8}$ ) is shown in red dashed line. (B) Venn diagram showing the overlap between 95% credible sets from mJAM-Forward, mJAM-SuSiE and MsCAVIAR.*

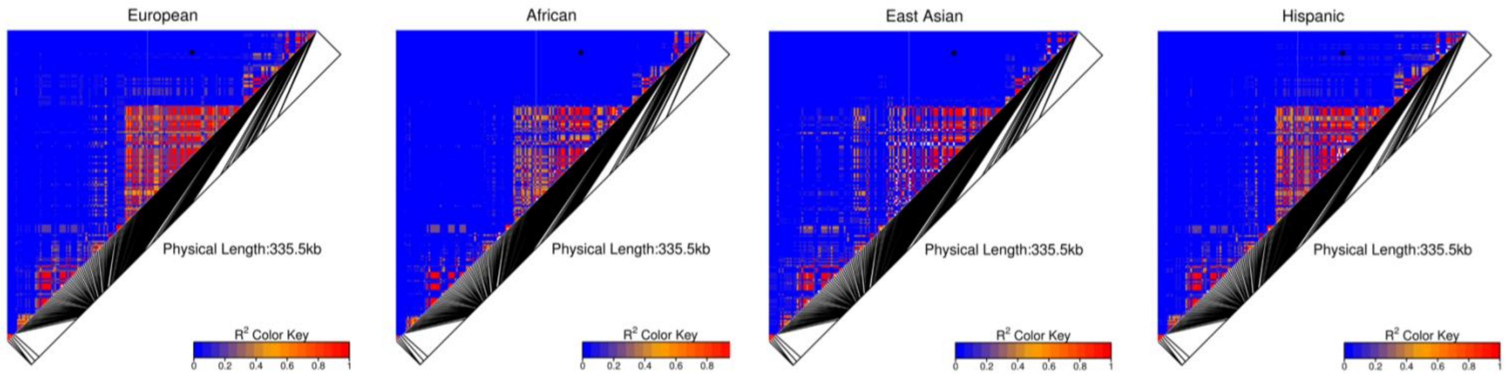


**Figure S 11 Ethnic-specific LD structure for 416 SNPs at chromosome 11 from position 101601661 to 103201661.**

*Analysis restricted to SNPs with meta-analyzed P-value < 0.001 and MAF > 2%. The pairwise correlation between 11:102440927:A:G and 11:102401661:C:T is marked in black asterisk.*

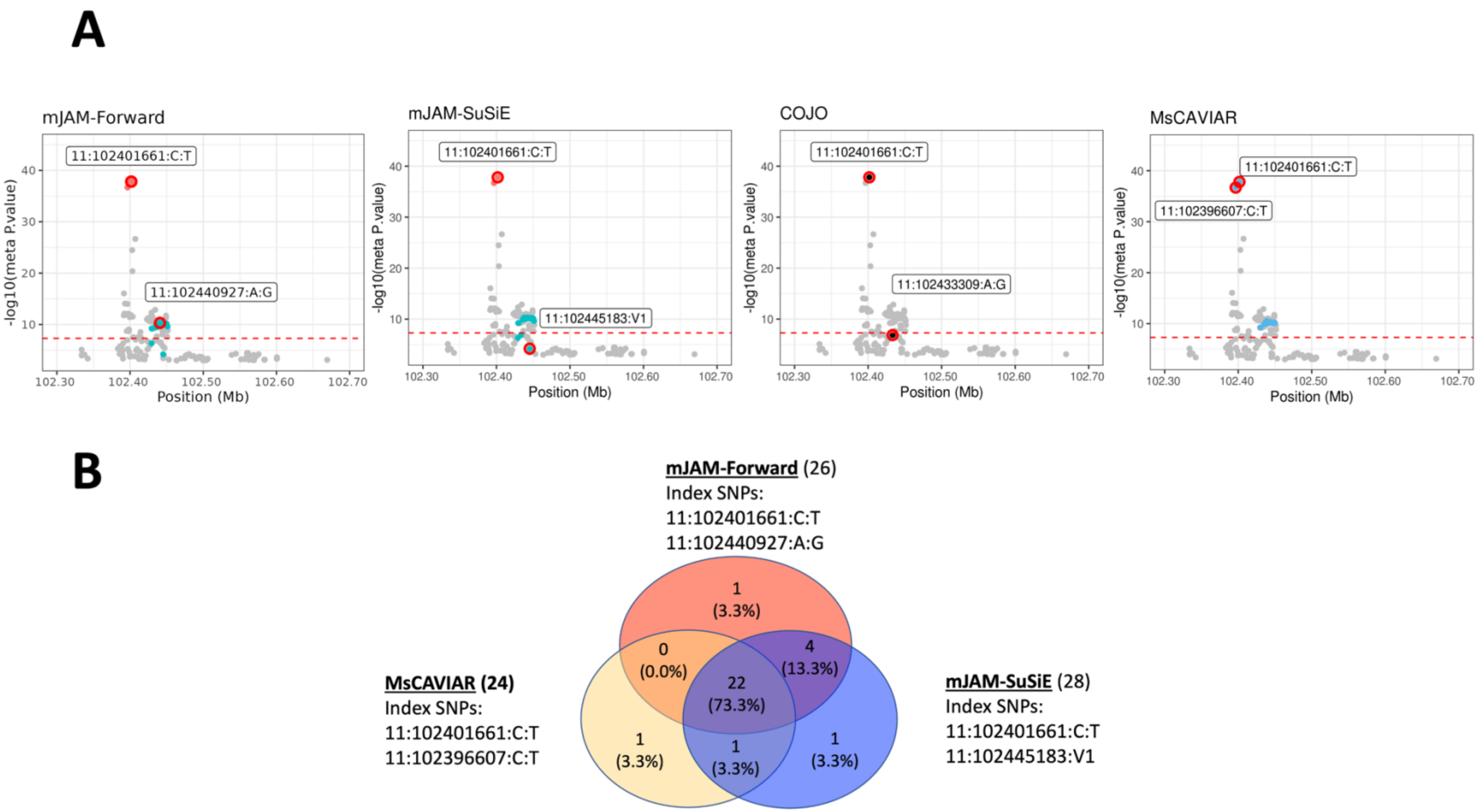


**Figure S 12 Fine-mapping results for chromosome 11 region from position 101601661 to 103201661.**

*(A) From left to right is mJAM-Forward 95% credible set, mJAM-SuSiE 95% credible set, index SNP selected by COJO, and MsCAVIAR 95% credible set. Index SNPs are circled in red. SNPs in the same credible sets are highlighted in the same color. Genome-wide significance ( $5 \times 10^{-8}$ ) is shown in red dashed line. (B) Venn diagram showing the overlap between 95% credible sets from mJAM-Forward, mJAM-SuSiE and MsCAVIAR.*

**Table S 1 Minimum, mean and median pairwise r2 of credible set SNPs with corresponding index SNP for the chr 12 region.**

*EUR, European; AFR, African American; HISP, Hispanic; ASI, East Asian.*

| | | EUR | AFR | HISP | ASI |
|---|---|---|---|---|---|
| mJAM-Forward | min | 0.62 | 0.35 | 0.74 | 0.33 |
| | mean | 0.98 | 0.95 | 0.98 | 0.94 |
| | median | 1.00 | 0.98 | 0.99 | 1.00 |
| mJAM-SuSiE | min | 0.60 | 0.58 | 0.62 | 0.41 |
| | mean | 0.97 | 0.95 | 0.97 | 0.95 |
| | median | 1.00 | 0.99 | 1.00 | 1.00 |
| MsCAVIAR | min | 0.62 | 0.35 | 0.74 | 0.33 |
| | mean | 0.98 | 0.95 | 0.98 | 0.94 |
| | median | 1.00 | 0.97 | 0.99 | 1.00 |

**Table S 2 Minimum, mean and median pairwise r2 of credible set SNPs with corresponding index SNP for the chr 10 region.**

*EUR, European; AFR, African American; HISP, Hispanic; ASI, East Asian.*

| | | EUR | AFR | HISP | ASI |
|---|---|---|---|---|---|
| mJAM-Forward | min | 0.95 | 0.81 | 0.95 | 0.97 |
| | mean | 0.96 | 0.98 | 0.99 | 1.00 |
| | median | 0.96 | 0.98 | 0.98 | 1.00 |
| mJAM-SuSiE | min | 0.33 | 0.12 | 0.13 | 0.24 |
| | mean | 0.79 | 0.69 | 0.73 | 0.72 |
| | median | 0.97 | 0.88 | 0.97 | 0.99 |
| MsCAVIAR | min | 0.95 | 0.81 | 0.97 | 0.98 |
| | mean | 0.96 | 0.98 | 0.99 | 1.00 |

| | median | 0.96 | 0.98 | 0.98 | 1.00 |
|---|---|---|---|---|---|

**Table S 3 Minimum, mean and median pairwise r2 of credible set SNPs with corresponding index SNP for the chr 11 region.**

*EUR, European; AFR, African American; HISP, Hispanic; ASI, East Asian.*

| | | EUR | AFR | HISP | ASI |
|---|---|---|---|---|---|
| mJAM-Forward | min | 0.35 | 0.16 | 0.38 | 0.35 |
| | mean | 0.96 | 0.91 | 0.96 | 0.95 |
| | median | 1.00 | 0.99 | 1.00 | 1.00 |
| mJAM-SuSiE | min | 0.22 | 0.07 | 0.22 | 0.13 |
| | mean | 0.42 | 0.23 | 0.44 | 0.41 |
| | median | 0.35 | 0.16 | 0.38 | 0.35 |
| MsCAVIAR | min | 1.E-05 | 3.E-03 | 7.E-03 | 9.E-02 |
| | mean | 6.E-02 | 7.E-02 | 7.E-02 | 2.E-01 |
| | median | 3.E-05 | 2.E-02 | 9.E-03 | 1.E-01 |
